# Dopamine transporter and synaptic vesicle sorting defects initiate auxilin-linked Parkinson’s disease

**DOI:** 10.1101/2022.02.04.479203

**Authors:** D J Vidyadhara, Mahalakshmi Somayaji, Nigel Wade, Betül Yücel, Helen Zhao, N Shashaank, Joseph Ribaudo, Jyoti Gupta, TuKiet T. Lam, Dalibor Sames, Lois E. Greene, David L. Sulzer, Sreeganga S. Chandra

## Abstract

Auxilin participates in the uncoating of clathrin-coated vesicles (CCVs), thereby facilitating synaptic vesicle (SV) regeneration at presynaptic sites. Auxilin (*DNAJC6*/*PARK19*) loss-of-function mutations cause early-onset Parkinson’s disease (PD). Here, we utilized auxilin-knockout (KO) mice to elucidate the mechanisms through which auxilin deficiency and clathrin-uncoating deficits lead to PD. We demonstrate that auxilin KO mice display the cardinal features of PD, including progressive motor deficits, α-synuclein pathology, nigral dopaminergic loss, and neuroinflammation. Through unbiased proteomic and neurochemical analyses, we demonstrate that dopamine homeostasis is disrupted in auxilin KO brains, including via slower dopamine reuptake kinetics *in vivo*, an effect associated with dopamine transporter misrouting into axonal membrane deformities in the dorsal striatum. We also show that elevated macroautophagy and defective SV protein sorting contribute to ineffective dopamine sequestration and homeostasis, ultimately leading to neurodegeneration. This study advances our knowledge of how presynaptic endocytosis deficits lead to dopaminergic vulnerability and pathogenesis of PD.

## INTRODUCTION

Presynaptic boutons of dopaminergic (DA) nigrostriatal neurons are sites of initiation for neurodegeneration in PD (Kordower et al., 2013). Nigrostriatal DA neurons possess long, hyperbranched axons as well as tonic firing properties that make them reliant on efficient synaptic vesicle (SV) recycling to maintain a steady state SV pool for neurotransmission (Vidyadhara et al., 2019). In presynaptic sites, SV recycling is supported by several endocytic pathways, including clathrin mediated endocytosis (CME), ultra-fast endocytosis, and bulk endocytosis (Chanaday et al., 2019). Clathrin coated vesicles (CCVs) are a common intermediate of these pathways. CCVs are uncoated by the coordinated action of auxilin (*DNAJC6*) or its ubiquitous homolog cyclin G-associated kinase (*GAK*), synaptojanin-1 (*SNJ1*), and endophilin-A1 (*ENDOA1*) with the chaperone Hsc70. Interestingly, mutations in all four genes (*DNAJC6*, *GAK*, *SNJ1*, *ENDOA1*) have been identified as causal or risk alleles for PD/parkinsonism, suggesting a major role for altered clathrin uncoating in the initiation of DA presynaptic site degeneration and pathogenesis of both familial and sporadic PD (Vidyadhara et al., 2019). Animal models carrying these mutations also show clathrin uncoating and presynaptic endocytosis defects (Cao et al., 2017; Song et al., 2017; Yoshida et al., 2018), however, how these disturbances result in characteristics of PD is unclear.

Auxilin is a brain-specific heat shock protein 40 family co-chaperone that functions to uncoat CCVs to nascent SVs by recruiting Hsc70 (Fotin et al., 2004; Ungewickell et al., 1995). Unlike other co-chaperones, auxilin has a limited number of substrates (Roosen et al., 2021) and only one described function. Loss-of-function, autosomal recessive mutations of the auxilin gene (*PARK19*) cause juvenile, early-onset PD (Edvardson et al., 2012; Elsayed et al., 2016; Köroğlu et al., 2013; Mittal, 2020; Ng et al., 2020; Olgiati et al., 2016; Ray et al., 2021). A recent study also shows *PARK19* mutations in late-onset PD (Gialluisi et al., 2021). *In vivo* presynaptic dopamine transporter (DAT) imaging on a *PARK19* patient revealed DA terminal loss, which supports a parkinsonism diagnosis and suggests that clathrin uncoating deficits impact DA presynaptic sites (Ng et al., 2020). Furthermore, LRRK2 mutations, a common genetic cause for PD, may exert some of its pathological actions through auxilin. In LRRK2 patient induced pluripotent stem cell derived DA neurons, LRRK2 phosphorylation of auxilin led to decreased auxilin levels and clathrin binding, resulting in accumulation of oxidized-dopamine and α-synuclein overexpression (Nguyen and Krainc, 2018). Whether loss-of-function mutations in auxilin can also trigger PD through these mechanisms is unknown. Nonetheless, these newfound links between auxilin and LRRK2 implicate a role for auxilin in both familial and sporadic PD. The relevance of auxilin to all forms of PD is underscored by the finding that GAK (*DNAJC26*) is a risk allele for sporadic PD (Nalls et al., 2014).

Prior to the discovery of auxilin *PARK19* mutations, auxilin knockout (KO) mice were generated and characterized for CME deficits (Yim et al., 2010). Analysis of synapses in deep cerebellar nuclei of adult auxilin KO mice revealed accumulation of CCVs and empty clathrin cages (lacking SV membrane). Similar ultrastructural alterations were seen *in vitro* in primary cortical and hippocampal neurons and were accompanied by defective SV endocytosis (Yim et al., 2010). These findings confirmed that auxilin functions in clathrin uncoating *in vivo*. To determine how a primary deficit in clathrin uncoating that occurs in all neuronal types lead to selective vulnerability of DA neurons in PD, we characterized auxilin KO mice for age-dependent nigrostriatal degenerative changes and investigated the underlying mechanisms. Here, we demonstrate that cytoplasmic dopamine accumulation, DAT mis-trafficking, SV sorting deficits and autophagic overload in dorsal striatal DA presynaptic sites of auxilin KO mice initiate behavioral and histochemical signatures of PD.

## RESULTS

### Auxilin KO mice develop age-dependent PD-like behavioral abnormalities

We performed a battery of behavioral assays to evaluate if auxilin KO mice develop age-dependent motor behavior abnormalities akin to PD patients. We monitored cohorts of wildtype (WT, C57BL/6J) and auxilin KO (congenic B6.-*Dnajc6^tm1Legr^*) mice longitudinally, assessing behavior at 3, 6, 9, 12, and 15 months of age. Locomotion and ambulatory behaviors were evaluated by the open field test. Auxilin KO mice behave like WT mice at 3 months of age but show a significant age-dependent decrease in overall distance travelled, starting at 9 months (Figure. 1a, b). Next, we tested the same cohorts on the balance beam test to evaluate motor coordination. We assessed the ability of mice to traverse a raised narrow beam by measuring the time taken to cross (Figure. 1c) and the number of runs performed in 1 minute (Figure. 1d). The performance of auxilin KOs was comparable to WT controls at 3 months (Figure. 1c, d), whereas it deteriorated in auxilin KOs at a later age with a significant deficit emerging at 9 months (Figure. 1c, d, Video 1). These results suggest that auxilin KOs appear to be normal at 3 months, whereas they become symptomatic by 9 months, exhibiting a progressive decrement in motor function at later ages. There was no difference in body weight between genotypes (Supplementary Figure. 1g). Furthermore, performance on the Rotarod and grip strength in auxilin KOs were comparable to that of WT (Supplementary Figure. 1a, e, f). In addition, auxilin KO mice do not exhibit anxiety-like behavior as evaluated by time spent in the inner and outer circle of an open field apparatus (Supplementary Figure. 1b, c), and by fecal pellet expulsion (Supplementary Figure. 1d). Together, these observations suggest that auxilin KO mice develop age-dependent, progressive motor deficits, consistent with *PARK19* and PD patients.

**Figure. 1:**
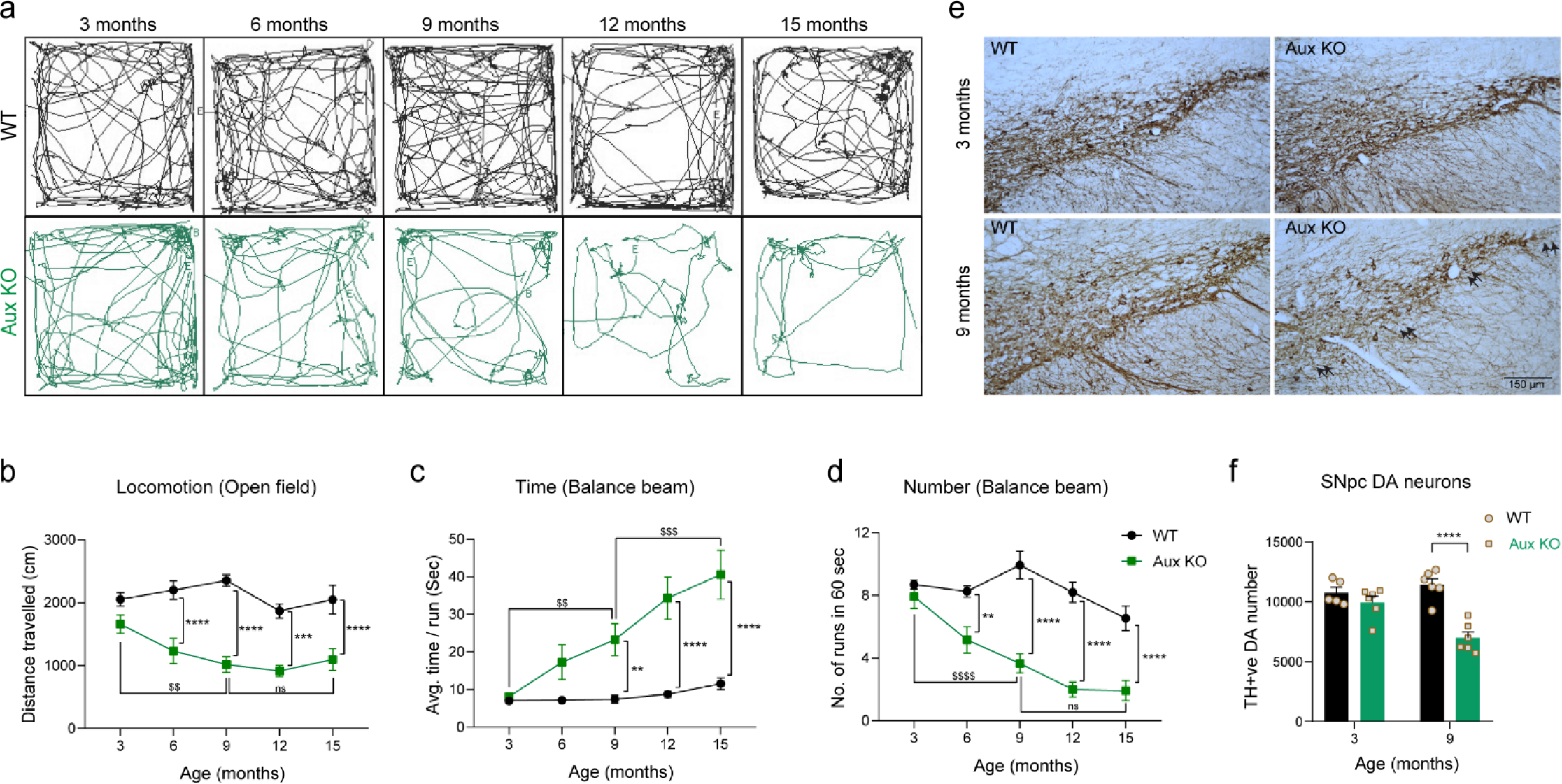
Auxilin KO mice develop progressive motor behavior deficits that are accompanied by nigral DA neuronal loss. **a**. Longitudinal open field locomotory behavior tracings of WT and auxilin KO mice (Aux KO) from 3 to 15 months of age, performed at 3 months interval. **b**. Distance travelled in open field as a function of age. At 3 months, Aux KOs (n=12) were comparable to WT (n=16). A progressive diminishment in locomotion is seen in Aux KOs with age, which was significant beyond 9 months compared to their performance at 3 months. **c.** Time taken to traverse balance beam. Aux KOs take longer to cross the beam with age, with a significant difference after 9 months of age. **d.** Number of runs performed in a minute on a balance beam. **e.** Representative images showing TH+ve SNpc DA neurons in WT and Aux KO midbrain sections at 3 and 9 months of age. Fewer DA neurons (arrows) were present in the SNpc of Aux KO mice at 9 months. Scale bar: 150 μm. **F.** Unbiased stereological counting of SNpc DA neurons. Note a significant (∼40%) loss of DA neurons is seen in 9-month-old Aux KO mice (n=5- 6/genotype). Statistics: For behavior, two-way repeated measure ANOVA followed by Sidak’s multiple comparison test was used. For stereology, Student’s t-test with Welch’s correction was used. *\*\*p<0.01, ***p<0.001, ****p<0.0001, ^$$^p<0.01, ^$$$$^p<0.0001, ^$$^p<0.01, ^$$$$^p<0.0001, ns= not significant*.

### Aged auxilin KO mice faithfully replicate cardinal histopathological signatures of PD

Motor symptoms in PD manifest due to the degeneration of DA neurons in the substantia nigra pars compacta (SNpc) when DA loss reaches a threshold of 40-50% (Poewe et al., 2017). We performed stereological quantitation of SNpc DA neurons, that are immunoreactive to tyrosine hydroxylase (TH), in WT and auxilin KOs to understand the cellular basis for the motor symptoms we observed. No change in DA neuron numbers was seen in auxilin KO mice at 3 months (Figure. 1e, f). However, at the symptomatic age of 9 months, a significant loss of DA neurons was observed (∼40%) (Figure. 1e, f), which did not increase further at 15 months (Supplementary Figure. 1h, i). Neuronal loss was distributed throughout the SNpc (Figure. 1e, arrows), as observed in models of α-synuclein overexpression (Chen et al., 2015) and vesicular dopamine storage deficits (Caudle et al., 2007), but unlike the ventrolateral loss seen in neurotoxic models (Vidyadhara et al., 2017). In addition, loss of DA neurons appears to be restricted to SNpc, as ventral tegmental area (VTA) TH expression and TH+ve neuron numbers were unchanged with age (Supplementary Figure. 2a-c), as in PD patients.

**Figure. 2:**
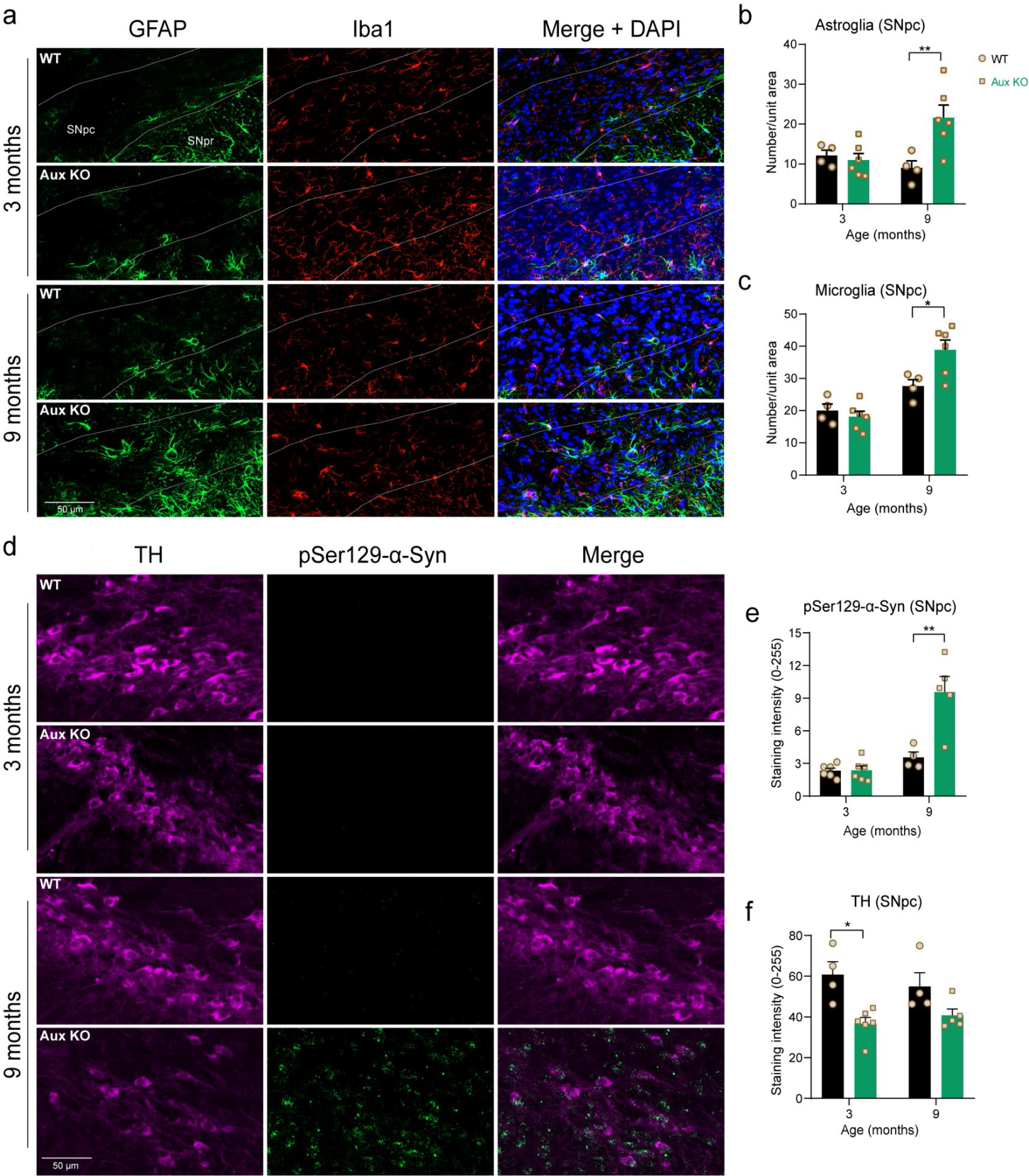
Aged auxilin KO mice exhibit gliosis and α-synuclein pathology. **a.** Representative images of SNpc and SN pars reticulata (SNpr) of WT and Aux KO mice at 3 and 9 months (n=5- 6/genotype) immunostained for the astroglial marker GFAP (green) and microglial marker Iba1 (red). Dashed line demarcates SNpc from SNpr. Scale bar: 50 μm **b.** Quantitation of GFAP+ve cells, show a significant astrogliosis at 9 months, but not at 3 months of age in the SNpc. **c.** Quantitation of Iba1+ve cells, show microgliosis in the SNpc of Aux KOs at 9 months. **d.** Representative images of the SNpc immunostained for pSer129-α-synuclein (green), a marker of α-synuclein pathology, co-stained with DA marker TH (magenta). Scale bar: 50 μm. **e.** Quantitation of pSer129-α-synuclein +ve punctate aggregates in SNpc, which showed an increase at 9-month-old Aux KOs, but not at 3 months. **f.** Quantitation of TH staining intensity in SNpc, which showed a moderate decrease at 3 months in auxilin KO mice but not at 9 months, suggesting retention of TH phenotype in the surviving neurons. Statistics: Student’s t-test with Welch’s correction. *\*p<0.05, **p<0.01*.

To assess if neurodegeneration was accompanied by neuroinflammation, we immunostained for glial fibrillary acidic protein (GFAP), an astroglial marker, and ionized calcium-binding adapter molecule 1 (Iba1), a microglial marker (Figure. 2a). At 3 months, the number of astroglia and microglia in the SNpc of auxilin KOs were comparable to WT, whereas at 9 months, significant astrogliosis and microgliosis was seen in auxilin KO mice (Figure. 2a-c).

Next, we tested if auxilin KO brains exhibit α-synuclein pathology, a hallmark of PD (Poewe et al., 2017). Strikingly, auxilin KO brains showed age-dependent α-synuclein pathology. Phosphorylated and aggregated α-synuclein as determined by pSer129-α-synuclein immunostaining was seen in the TH+ve SNpc at 9 months, but not at 3 months of age in auxilin KOs (Figure. 2d, e). Immunostaining also revealed a moderate decrease in TH expression at 3 months, with no significant change at 9 months of age in auxilin KOs (Figure. 2d, f). The pSer129- α-synuclein pathology was also seen in the VTA at 9 months of age, but the expression did not reach significance (Supplementary Figure. 2a, c). These compelling data demonstrate that auxilin KOs develop typical age-related parkinsonian pathology. Auxilin KO mice are thus a reliable and robust model for PD.

### Proteomic analysis of auxilin KO mice brains implicate defective dopamine degradation

To gain unbiased insights into the consequences of auxilin loss-of-function, we performed proteomic analysis on whole brain and synaptosomes fractions from 3-month-old, WT and auxilin KO mice by label-free quantification mass spectrometry (LFQ-MS). WT and auxilin KO brains (n=3/genotype) were analyzed in technical triplicates. We detected a total of 2851 proteins in the whole brain proteome, 22 of which were significantly changed in KO samples (Figure. 3a, b; Supplementary Table 1). We observed an expected decrease in auxilin levels and a compensatory increase in the auxilin homolog GAK, as previously published (Yim et al., 2010). Many of the prominent proteins whose levels are changed are linked to PD and neurodegeneration, including RAB3B, TBCD, ACAP2, HEBP1, WDFY1 and NNTM which are decreased, while CRYAB, PRIO, and NMRL1 are increased in auxilin KO brains (Figure. 3a, b; Supplementary Table 1). Notably, we did not identify any altered Golgi resident or trafficking proteins in auxilin KO brains. Ingenuity Pathway Analysis (IPA) revealed that the top pathways were highly overlapping and involve in the degradation of lysine and degradation of choline and monoaminergic neurotransmitters, including dopamine (Figure. 3c, d). Interestingly, a decrease in AL7A1 appears to drive the top canonical pathways (Figure. 3c). AL7A1 or aldehyde dehydrogenase 7A1 (ALDH7A1) is a multifunctional enzyme which plays crucial role in detoxification of reactive aldehydes and oxygen species that are generated during monoaminergic neurotransmitter metabolism (Brocker et al., 2011). Aldehydes that accumulate due to ALDH7A1 loss-of-function hinder dopamine synthesis (Clayton, 2020).

**Figure 3:**
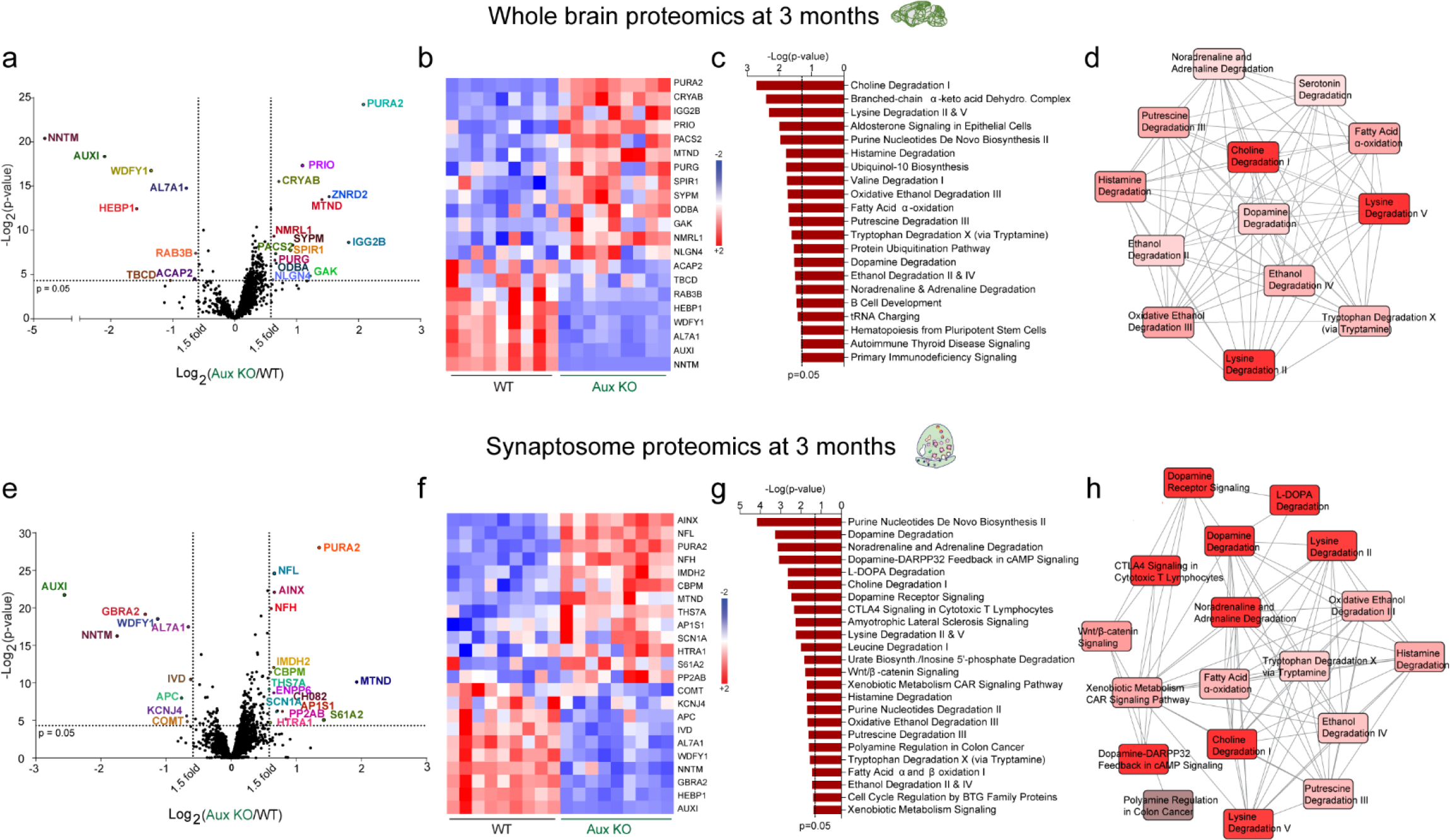
Whole brain and synaptosome proteomics reveal dopamine degradation dysfunction in young auxilin KOs. **a.** Volcano plot of whole brain proteome of Aux KOs compared to WT (age=3 months, n=3 mice/genotype). Proteins that exhibit a 1.5-fold change (vertical dotted lines) and a p-value of 0.05 (Student’s t-test) or lesser (horizontal dotted line) were considered as significantly changed. Among the 22 proteins that significantly changed, 8 were decreased (left quadrant) and 14 were increased (right quadrant). **b.** Heat map of significantly changed proteins in whole brain homogenates of WT and Aux KOs for all 9 technical replicates (3 technical replicates/mouse). Red indicates an increase (+2) and blue indicates decreased levels (-2). **c.** Pathways that are significantly (p<0.05) affected in whole brain of Aux KOs as determined by IPA. **d.** Diagram showing the overlap of significantly affected pathways, where intense red depicts most affected and light red depicts moderately affected pathways. **e.** Volcano plot of synaptosome proteome of Aux KOs compared to WT (age=3 months, n=3 mice/genotype). Out of 24 proteins that were significantly changed (1.5-fold, p<0.05, Student’s t-test), 9 were decreased (left quadrant) and 15 were increased (right quadrant). **f.** Heat map of significantly changed proteins in synaptosome preparations from Aux KOs in comparison to WT for each technical replicate. Red indicates an increase (+2) and blue indicates decreased levels (-2). **g.** Significantly affected pathways due to synaptosome proteomic changes as determined by IPA. **h.** Overlap of significantly affected pathways showing highly affected (intense red) and moderately affected (light red) pathways.

Proteomic analysis of synaptosomal fractions identified 3124 proteins, 24 of which were significantly dysregulated (Figure. 3e, f; Supplementary Table 2). Along with the expected downregulation of auxilin, KO mice showed decreased AL7A1, NNTM and WDFY1, and an upregulation in PURA2 and MTND, proteins which were also significantly changed in whole brain proteomic experiments (Figure. 3a, b). Three neurofilaments NFL, NFH, and AINX were upregulated and are candidate biomarkers for axonal damage in PD (Bäckström et al., 2020). A crucial dopamine metabolizing enzyme COMT (Myöhänen et al., 2010) (catechol-o- methyltransferase) was also significantly decreased in synaptosome preparations of auxilin KOs. HTRA1, PP2A, KCNJ4 and APC were a few PD-linked proteins that were also dysregulated (Figure. 3e, f; Supplementary table 2). In all, we find alterations in a high number of proteins linked to PD (13/23), including key dopamine metabolism enzymes. As the synaptosome preparations were purified from whole brain, these findings strongly suggests that loss-of-auxilin selectively impacts DA neurons. This is also evident from the IPA analysis, where the top affected pathways are related to dopamine degradation (Figure. 3g, h). A high fraction of the canonical pathways predicted for the whole brain analysis were replicated in IPA analysis for the synaptosome samples (10/21, compare Figure. 3g, h with 3c, d), suggesting a major impact on the function of DA synapses upon loss-of-auxilin.

To evaluate if the disruption in dopamine degradation predicted by the proteomic analysis of young auxilin KO brains (Figure. 3) leads to activation of downstream neurodegenerative pathways at an older age, we performed LFQ-MS on synaptosomes from 9-month-old auxilin KO mice. In this preparation, we still observed a compensatory increase in GAK in auxilin KO brains. NFL, AINX, and CADH2 were also upregulated as in the 3-month data set, reinforcing their potential as biofluid biomarkers of PD (Bäckström et al., 2020) (Supplementary Figure. 3a, b). Along with the expected downregulation of auxilin, mTOR, a key cell survival protein and an autophagy regulator which helps maintain striatal DA projections (Kosillo et al., 2019) and linked to PD (Querfurth and Lee, 2021), was decreased in auxilin KO mice. RGS6, a critical regulator of dopamine feedback signaling in nigrostriatal DA neurons and a modulator of PD pathology (Ahlers-Dannen et al., 2020) was also downregulated (Supplementary Figure. 3a, b; Supplementary table 4). IPA revealed highly overlapping autophagic pathways such as ILK, P13K/AKT, mTOR, and AMPK signaling, along with oxidative stress, DA signaling, and ubiquitination pathway as dysregulated in auxilin KOs at 9 months (Supplementary Figure. 3c), which are directly linked to neurodegeneration in PD (Poewe et al., 2017). Together, our proteomic analyses suggests that dysfunction of dopamine homeostasis in presynaptic sites is likely to be an early pathogenic event in auxilin-linked PD.

### Disrupted striatal dopamine homeostasis in auxilin KO mice

To directly monitor dopamine homeostasis, we measured the levels of dopamine and its metabolites in dorsal striatum of WT and auxilin KO mice using high performance liquid chromatography (HPLC). Dopamine levels were moderately decreased (14.5%) at 3 months in auxilin KO mice compared to WT controls, whereas loss was more pronounced at 9 months of age (52%, Figure. 4a) when motor deficits are seen (Figure. 1). We also measured serotonin levels, which were unchanged in auxilin KOs (Supplementary Figure. 4a). Next, we evaluated the levels of dopamine metabolites 3,4- dihydroxyphenylacetic acid (DOPAC) and homovanillic acid (HVA), which are intra- and extra- cellular metabolites, respectively (Figure. 4e). Interestingly, DOPAC levels were significantly increased at 3 months (42%, Figure. 4b), even though dopamine levels are modestly decreased. DOPAC is a catabolite of cytosolic (non-vesicular) dopamine (Figure. 4e). Upregulation of DOPAC suggests cytosolic dopamine accumulation (Karoum et al., 1994), which is known to be toxic, leading to oxidative stress and proteostasis deficits which culminate in neurodegeneration in both familial and sporadic models of PD (Masato et al., 2019). 3-Methoxytyramine (3-MT) is a dopamine metabolite, formed by direct catabolism of unused dopamine in the synaptic-cleft by COMT (Myöhänen et al., 2010) (Figure. 4e). 3-MT levels were significantly lower in auxilin KOs (Figure. 4c) suggestive of a decrease in dopamine release (Waldmeier et al., 1981), and reflective of the downregulation of COMT seen in the proteomics data (Figure. 3e, f). Both DOPAC and 3- MT are metabolized further to HVA outside the DA termini (Figure. 4e), whose level did not change at 3 months in auxilin KOs (Figure. 4d). This is possibly due to a balancing out of HVA levels attained by increased DOPAC and decreased 3-MT levels. At 9 months when motor abnormalities are apparent, both 3-MT and HVA levels were significantly decreased in auxilin KO mice (Figure. 4c, d), which is also the case in PD patients. Indeed, decreased HVA levels have also been observed in the cerebrospinal fluid of patients with auxilin mutations (Ng et al., 2020). DOPAC levels were unchanged at 9 months. Levels of the serotonin metabolite 5- hydroxyindoleacetic acid (5-HIAA) did not change at 3 and 9 months (Supplementary Figure. 4b), suggesting dopamine-selective dysfunction. Overall, assessment of dopamine and its metabolites in dorsal striatum support the premise that dopamine homeostasis is altered in auxilin KO mice.

**Figure. 4:**
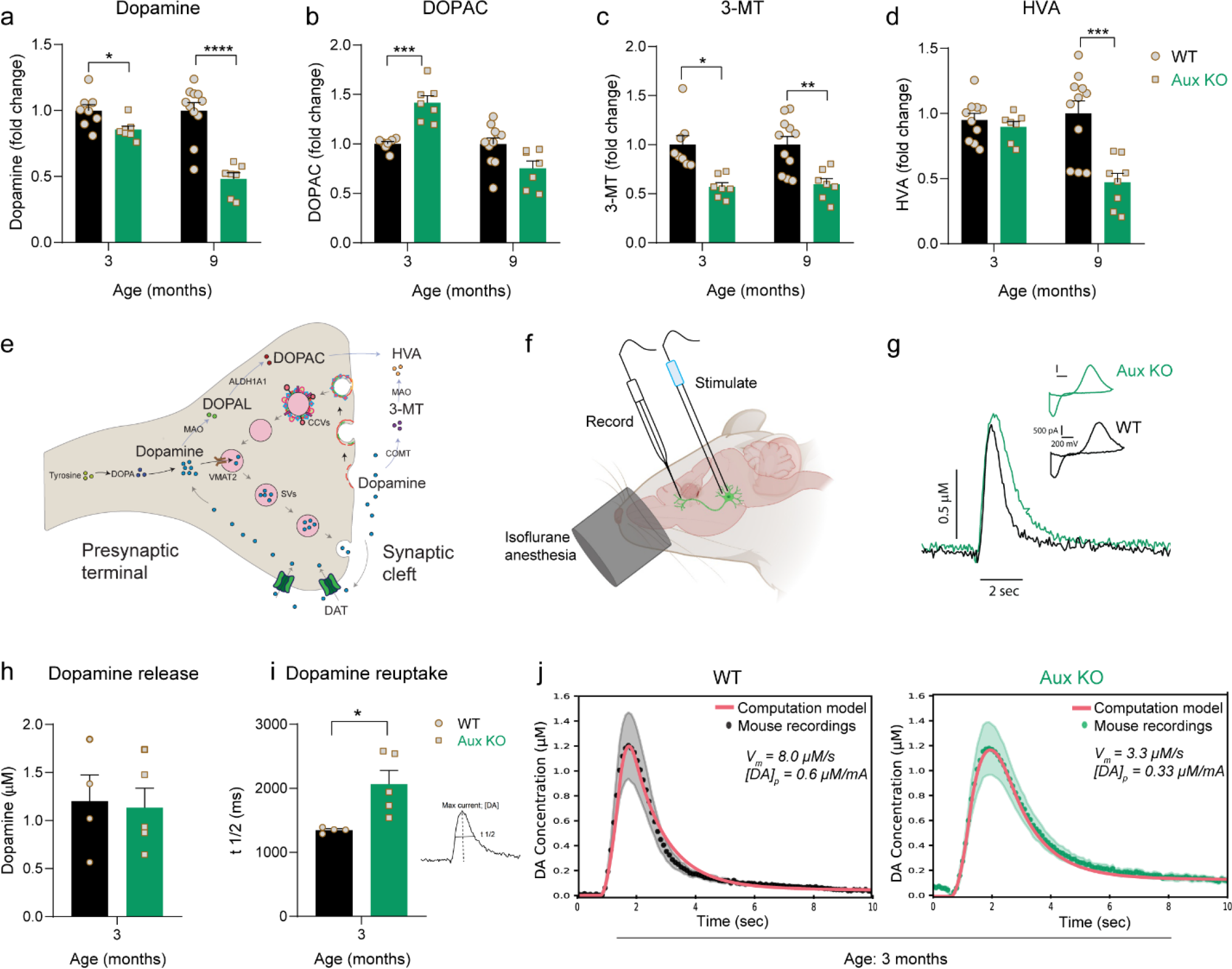
Dopamine catabolism and dopamine reuptake deficits in young auxilin KO mice. **a.** Dopamine levels in the dorsal striatum of WT and Aux KO mice. Note a modest decrease in dopamine at 3 months, but a larger decrease at 9 months of age (n=7-11 mice/genotype). **b.** DOPAC levels in the dorsal striatum. At 3 months, a significant increase in the intracellular dopamine catabolite DOPAC is seen in Aux KOs, whereas no change is observed at 9 months. **c.** 3-MT levels in the dorsal striatum. An extra-synaptic dopamine catabolite 3-MT was decrease both at 3 and 9 months of age in Aux KOs. **d.** HVA levels in the dorsal striatum. HVA, another extra- synaptic dopamine catabolite did not change at 3 months but decreased significantly at 9 months in Aux KO mice. **e**. Schematic showing compartmentalization of dopamine and its catabolites in intra- and extra-synaptic space. **f.** Schematic showing the location of FSCV recording electrode in the dorsal striatum and the bipolar stimulating electrode in the ventral midbrain of mice under isoflurane anesthesia. **g.** Example trace of evoked dopamine release following stimulation of midbrain DA neurons by 30 pulses at a constant 50-Hz frequency in 3-month-old Aux KO and WT mice (scale: y axis, 0.5 µM dopamine; x axis, 2 sec). **h.** Dopamine release in the dorsal striatum. No significant change in dopamine release was seen in Aux KO mice when compared to WT (n=4-5/genotype). **i.** Dopamine reuptake in the dorsal striatum. Reuptake kinetics measured by time taken to clear half the dopamine from its peak levels (t1/2) was significantly delayed in Aux KO mice. Statistics: Student’s t-test with Welch’s correction. **j.** Best-fits of computational model of dopamine (DA) release (red lines) to averaged FSCV recordings in the dorsal striatum of WT (Black dots; *R*^2^ = 0.98, n = 4) and Aux KO (Green dots; *R*^2^ = 0.99, n = 5) mice. Black/green ribbons report SEM. Dopamine release from Aux KO mice is closely fit by a ∼60% reduction in DAT activity (parameter *V*_*m*_) and a ∼45% reduction in dopamine release per electrical pulse (parameter [*DA*]_*p*_). *\*p<0.05, ***p<0.001, ****p<0.0001*.

### Dopamine reuptake is dysfunctional in young auxilin KO mice

Extracellular dopamine in the striatum is pumped back into DA axons by the dopamine transporter (DAT). Therefore, DAT controls the level of presynaptically available dopamine and is a key regulator of dopamine compartmentalization and homeostasis (Bu et al., 2021). DAT KO mice show a decreased striatal dopamine levels and release (Jones et al., 1998). We evaluated extracellular dopamine clearance on a sub-second timescale in the dorsal striatum of auxilin KOs using fast scan cyclic voltammetry (FSCV) *in vivo*. The SNpc was stimulated using a 30 pulses of 50-Hz stimuli (0.6 sec) paradigm that drives burst firing by nigral DA neurons. This causes dopamine build-up in the extracellular space at levels sufficient to saturate DAT and to be detected by the carbon fiber electrode placed in the dorsal striatum (Somayaji et al., 2020) (Figure. 4f, Supplementary Figure 5a). Figure. 4g and Supplementary Figure. 5b depict the time course of evoked dopamine release and its clearance in the dorsal striatum, along with the characteristic background-subtracted voltammogram at the maximum oxidation peak for WT and auxilin KO mice. Surprisingly, evoked dopamine release was not significantly different between WT and auxilin KOs (Figure. 4h). Interestingly, dopamine reuptake kinetics as measured by the time taken to clear 50% of the dopamine from its peak levels (t1/2) was significantly delayed in auxilin KO mice (Figure. 4i), suggesting a deficit in DAT function.

**Figure. 5:**
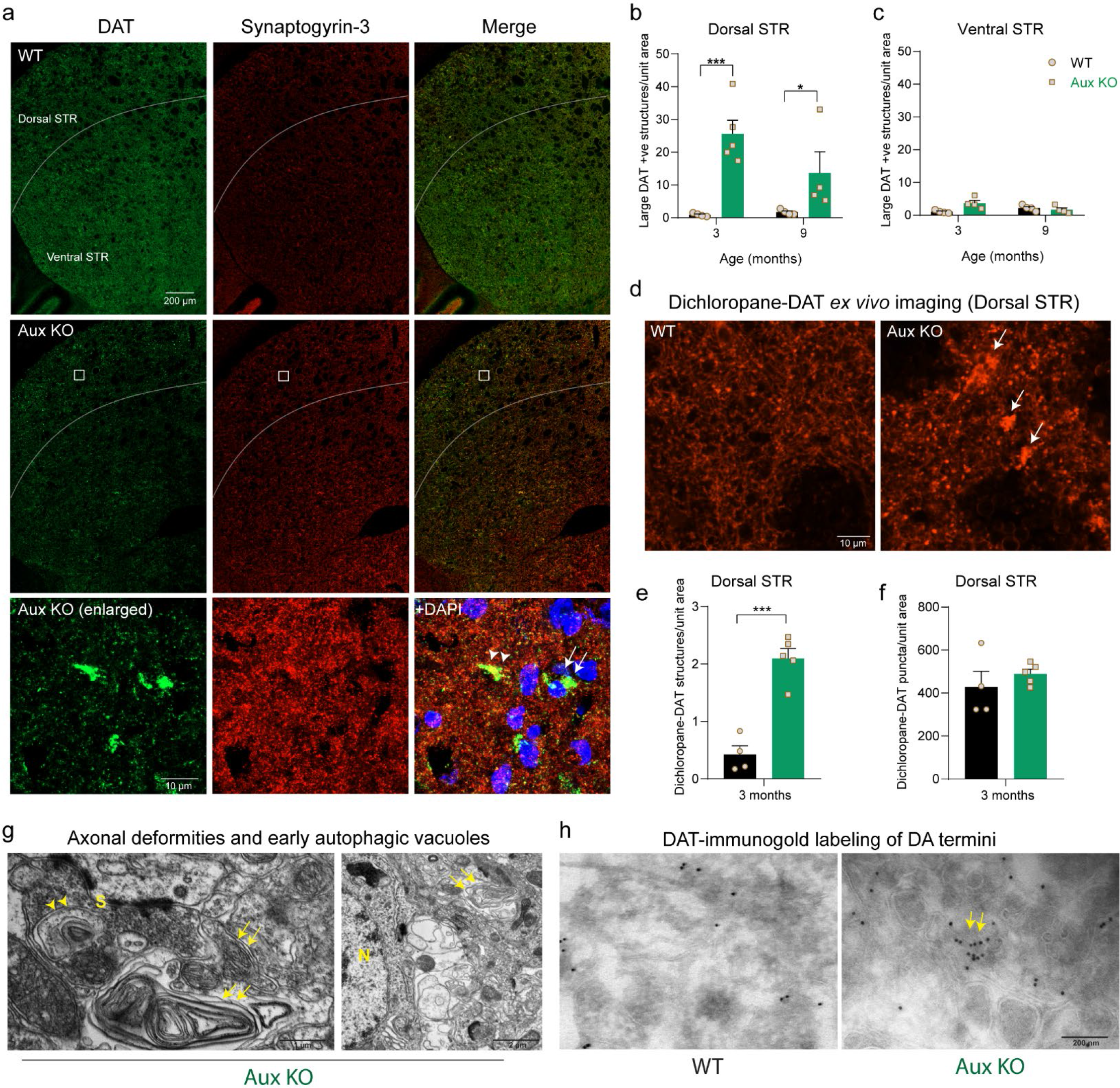
DAT+ve axonal deformities in the dorsal striatum of auxilin KOs. **a.** Representative images of dorsal and the ventral striatum (STR) in WT and Aux KO mice, immunostained for DAT and synaptogyrin-3. Note large DAT+ve structures in the dorsal STR of Aux KOs, that are absent in the ventral STR. Scale bar: 200 µm. These DAT+ve structures were juxtaposed to presynaptic sites as seen by colocalization with synaptogyrin-3 (Aux KO, enlarged, arrowhead), as well as in the soma marked by DAPI staining (Aux KO, enlarged, arrow). Scale bar: 10 µm. **b.** Number of DAT+ve structures/unit area in the dorsal striatum. Aux KO showed a significantly higher DAT+ve structures both at 3 and 9 months. **c.** Number of DAT+ve structures in the ventral striatum. DAT+ve structures were not observed in ventral striatum in Aux KOs. **d.** Representative images of *ex vivo* staining of dichloropane–rhodamine red-X in the dorsal striatum of WT and Aux KO mice. DA axonal projections and presynaptic sites appear as small puncta, whereas axonal deformities appear as large dichloropane-DAT+ve structures (arrows). Scale bar: 10 µm **e.** Number of large dichloropane-bound DAT+ve structures/unit area in the dorsal striatum, were significantly higher in Aux KOs. **f.** Number of small dichloropane-bound DAT+ve puncta, was not altered in Aux KO dorsal striatum in comparison to WT. **g.** EM of dorsal striatum of Aux KO mice showing large axonal whirl like deformities (arrows), which were present ubiquitously, closer to both synaptic terminals (S) and soma (N: nuclei). Early autophagic vacuole like structures were also seen in dorsal striatum (arrow heads), closer to axonal whirls. Scale bar: 1 µm and 2 µm. **h.** DAT- immunogold labeling of dorsal striatum that mark only DA axonal termini showed dispersed labeling in WT. In Aux KOs, DAT-immunogold clusters were seen in the dorsal striatum (arrows). Scale bar: 200 nm. Statistics: Student’s t-test with Welch’s correction. *\*\*\*p<0.001*.

To further analyze the difference in dopamine reuptake kinetics, we used a novel computational model of dopamine release derived from previous studies (Venton et al., 2003; Walters et al., 2014) to fit averaged FSCV traces from WT and auxilin KO mice. We found that the wider dopamine peak from auxilin KO mice can be closely fit by a ∼60% reduction in DAT activity (parameter *v*_*m*_) and a ∼45% reduction in dopamine release per electrical pulse (parameter *DA*_*p*_) compared to WT mice (*v*_*m*_ = 3.3 in auxilin KO vs 8.0 μM/s in WT, *DA*_*p*_ = 0.333 in auxilin KO vs 0.6 μM/mA in WT), while holding all other major parameters constant (Figure. 4j, Supplementary Figure. 5c). While DAT deficiency seen here was consistent with our FSCV recordings, the decrease in dopamine release in auxilin KOs in the computational model deviated from FSCV observations (Figure. 4h). However, decreased neurotransmitter release is expected in auxilin KOs as these mice have previously been shown to have SV recycling defects (Yim et al., 2010). Our neurochemical analyses show a decrease in the extracellular dopamine metabolite 3-MT (Figure. 4c) in the dorsal striatum of auxilin KO which also indicate dopamine release defects. Thus, it appears that a larger decrease in DAT activity is masking a decrease in dopamine release and can account for the minimal difference in total evoked dopamine release observed between WT and auxilin KO mice in the FSCV recordings.

### Auxilin KO mice exhibit DAT deformities in the dorsolateral striatum

To visualize DAT in auxilin KO mice, we performed immunohistochemistry for DAT, co-labeling with the presynaptic SV protein synaptogyrin-3 in dorsal striatum (Figure. 5a). We found large DAT+ve structures (6- 8 µm) in the dorsal striatum, but not in the ventral striatum of auxilin KO brains (Figure. 5a-c, Supplementary Figure. 6a), like structures that have been described in synaptojanin-1 knock-in mice (Cao et al., 2017). These structures were absent in WT, but ubiquitous in the dorsolateral striatum of auxilin KOs, localizing both with presynaptic sites (Figure. 5a, enlarged, arrowhead) and closer to the soma (as marked by DAPI) (Figure 5a, enlarged, arrows, Supplementary Figure. 6b). These DAT+ve structures were noted in auxilin KOs at both 3 and 9 months of age, though they were significantly higher (∼40%) at the earlier time point (Figure. 5b). Synaptogyrin-3, which marks presynaptic termini and a known interactor of DAT, did not exhibit a change in distribution or expression level (Figure. 5a, Supplementary Figure. 7f). Glutamatergic and GABAergic termini, which were imaged by staining for vesicular glutamate transporters (VGLUT1) and vesicular GABA transporter (VGAT), respectively, did not exhibit such structures (Supplementary Figure. 7a-c). As DAT is typically localized to DA axonal projections (Block et al., 2015), these observations suggest that the large DAT+ve structures seen at the dorsal striatum of auxilin KO may be DA axonal membrane deformities.

**Figure. 6:**
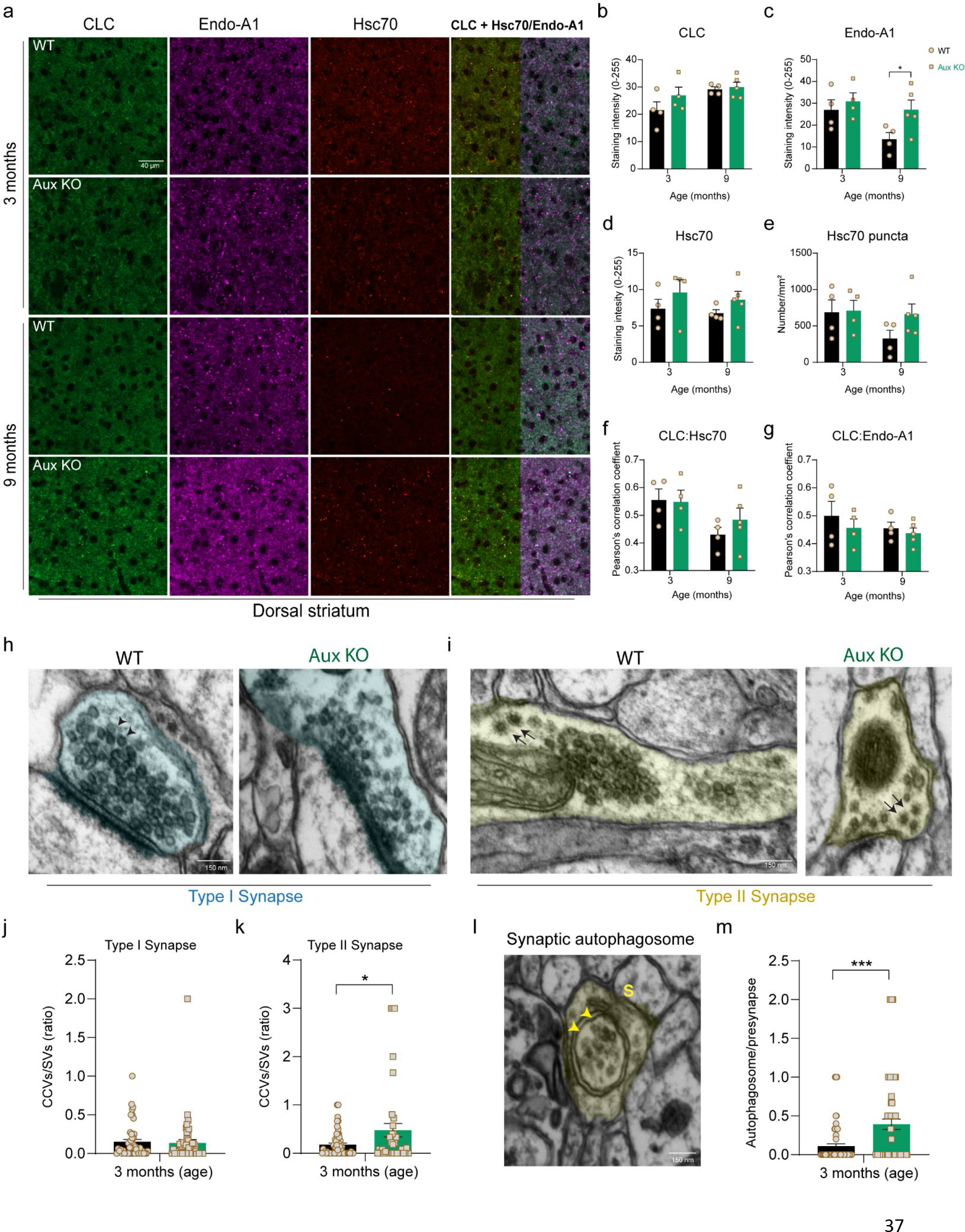
CCVs are cleared by synaptic autophagy in Auxilin KOs. **a.** Representative images of dorsal striatum immunostained for clathrin light chain (CLC), endophilin-A1 (Endo-A1) and Hsc70, in WT and Aux KO mice at 3 and 9 months of age. Scale bar: 40 µm. **b.** CLC expression the dorsal striatum, was not altered in Aux KOs. **c.** Endo-A1 expression in the dorsal striatum, was not changed at 3 months but was increased at 9 months in Aux KOs. **d.** Hsc70 expression in the dorsal striatum, which was also not altered**. e.** Number of Hsc70+ve puncta in the dorsal striatum is unaltered. **f.** Colocalization of Hsc70 with CLC in WT and Aux KOs. **g.** Endo-A1 colocalization with CLC. **h.** Representative EM image of a Type I excitatory presynapse with SVs (arrow heads) from dorsal striatum of WT and Aux KO. Scale bar: 150 µm. **i.** Representative EM image of a Type II inhibitory presynapse in the dorsal striatum of WT and Aux KO mice with SVs and CCVs (arrows). Note Aux KO presynapse showing a decrease accumulation of CCVs, accompanied by a decrease in SVs. Scale bar: 150 µm. **j.** The CCV to SV ratio in Type I synapses of dorsal striatum (Age: 3 months). This ratio was not altered in Aux KOs. **k.** The CCV to SV ratio in Type II synapses of dorsal striatum, which was significantly increased in Aux KOs in comparison to WT. **l.** EM image of a Type I synaptic terminal (S) in the dorsal striatum of Aux KOs showing double membraned autophagosomes containing CCVs (arrows). **m.** Number of autophagosomes per presynaptic terminal in the dorsal striatum, which was significantly increased in Aux KOs compared to WT (Age: 3 months). Statistics: Student’s t-test with Welch’s correction. *\*p<0.05, ***p<0.001*.

**Figure. 7:**
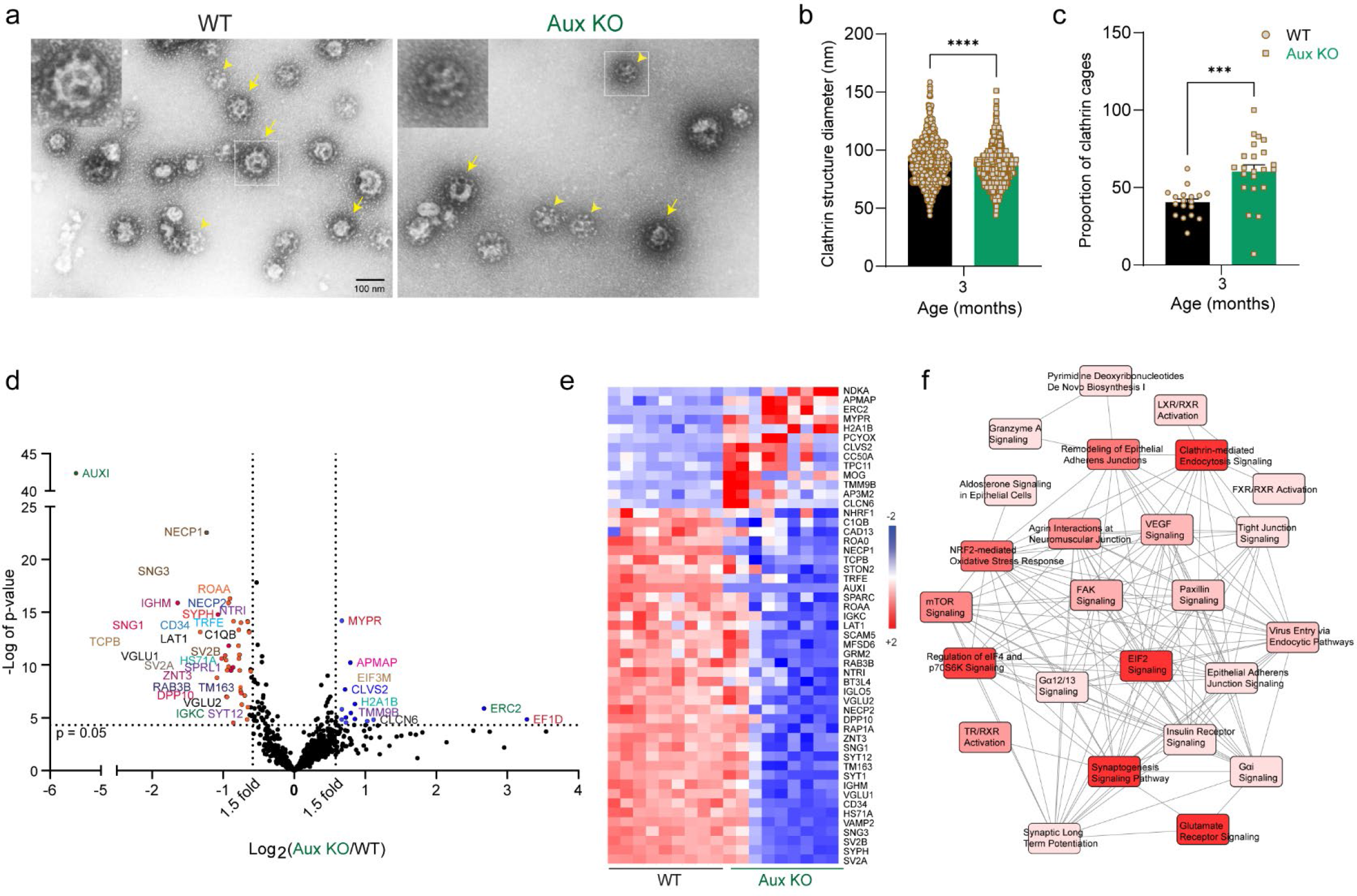
EM and proteomics of CCVs reveals SVs with variable membrane composition in auxilin KOs. **a.** Representative EM images of CCV preparation showing CCVs (arrows) and empty clathrin cages (arrow heads) in WT and Aux KO mice. Scale bar: 100 nm. **b.** Diameter of clathrin structures (CCVs + clathrin cages), which was significantly smaller in Aux KO samples compared to WT. **c.** Proportion of clathrin in WT and Aux KO mice. Note, Aux KOs show significantly larger proportion of clathrin cages. **d.** Volcano plot of CCV proteome of Aux KOs in comparison to WT (n=14 mice/experiment, N=3 experiments/genotype). Proteins that were changed 1.5-fold (vertical dotted lines) with a p-value of 0.05 (Student’s t-test) or lower (horizontal dotted line) were considered as significantly changed. Among the 49 proteins that were significantly changed, 37 were decreased and 12 were increased. **e.** Heat map of significantly changed proteins in Aux KOs in comparison to WT for each experiment (3 technical replicates per experiment). Red indicates increased expression (+2), and blue indicates a decrease (-2). **f.** Pathways that are significantly affected (p<0.05) in Aux KO mice due to CCV proteome changes, and their overlap. Pathways depicted in intense red are highly affected, whereas the light red are moderately affected. Statistics: Student’s t-test with Welch’s correction. *\*\*\*p<0.001, ****p<0.0001*.

To confirm that the DAT+ve structures are membrane-bound and surface accessible, we performed *ex vivo* imaging using the membrane DAT ligand dichloropane which binds to cell surface DAT preferentially from the extracellular side. Dichloropane was conjugated with rhodamine red-X, as described previously (Fiala et al., 2020) to obtain the dichloropane–rhodamine red-X probe (Fiala et al., 2020). Fresh striatal slices were incubated in artificial cerebrospinal fluid (ACSF) containing dichloropane probe (100 nM, 45 mins) and imaged for membrane-bound DAT. The number of small dicholoropane-DAT+ve punctum that represent DA terminal varicosities in the dorsal striatum were not altered in auxilin KOs at 3 months of age (Figure. 5d, f). However, auxilin KOs revealed large dicholoropane-DAT+ve structures (Figure. 5d, e, arrows) similar to those in the dorsal striatum of fixed brains, suggesting that the DAT structures are membrane accessible. Next, we performed an ultrastructure evaluation of dorsal striatum by electron microscopy (EM), which revealed multilayered axonal whirls in auxilin KO brains (Figure. 5g, Supplementary Figure. 8a, arrows). Additionally, there were early autophagic vacuole-like structures close to these axonal deformities (Figure. 5g, arrow heads). We performed immunogold labeling of DAT in the dorsal striatum, which revealed a uniform distribution of DAT-immunogold particles in WT, denoting DA axonal projections and presynaptic sites (Figure. 5h). In contrast, DAT-immunogold clusters were observed in auxilin KOs (Figure. 5h, arrows, Supplementary Figure. 8b, arrows), indicative of axonal membrane deformities of DA projections. Collectively, these observations suggest that DAT is mis-trafficked and trapped in large axonal deformities, which hinder DAT function in dopamine reuptake.

**Figure. 8:**
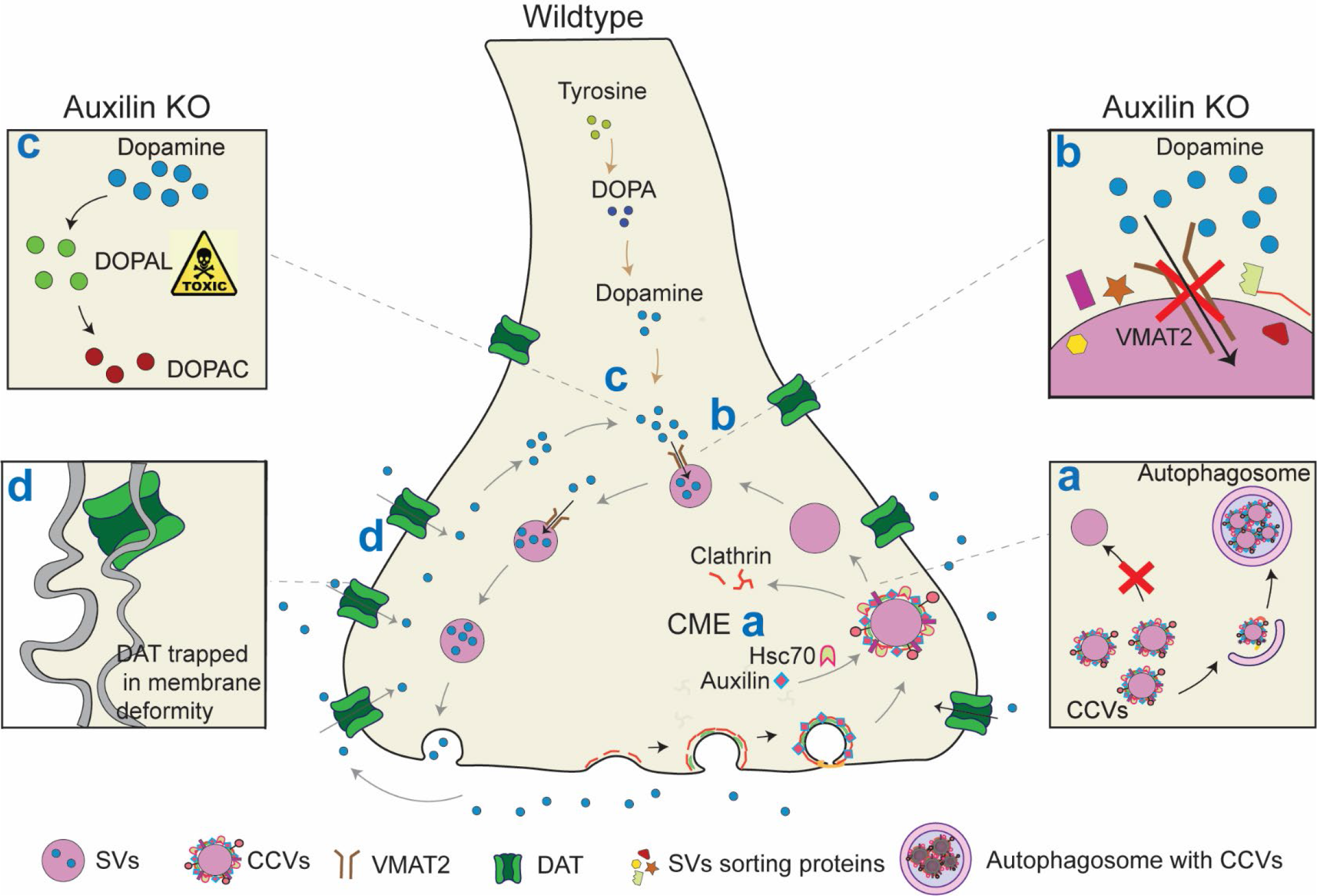
Schematic showing dopamine compartmentalization defects in a DA presynapse of auxilin KO mice. **a.** In a WT DA presynapse, after release of dopamine, SVs are recycled principally through CME. Clathrin forms a coat on the nascent SV membrane to form CCVs with the aid of adaptors. Auxilin recruits Hsc70 to CCVs and functions in its uncoating to generate SVs. In Aux KO mice, CCV uncoating is affected, leading to their accumulation and their subsequent clearance by an enhanced synaptic autophagy. Consequently, there is an imbalance in CCV to SV ratio. **b.** In WT DA synapses, dopamine once synthesized is immediately sequestered into the SVs by the vesicular transporter VMAT2. Aux KO mice have SVs with varied protein stoichiometry and a decrease in the copy number of vesicular transporters, hindering dopamine sequestration to SVs. **c.** In WT, there is minimal cytosolic dopamine present. In Aux KO, due to aforementioned events, there is an elevation in cytosolic dopamine, which is oxidized to its toxic intermediates such as DOPAL and DOPAC. **d.** Normally, DAT plays a pivotal role in dopamine reuptake and replenishing presynaptic vesicular dopamine for future release. In Aux KOs, DAT is misrouted and trapped in the axonal membrane whirls/deformities in the DA projections, compromising dopamine reuptake.

### Increased presynaptic autophagy in auxilin KO mice clear CCVs

We evaluated the distribution pattern of endocytic partners of auxilin in the dorsal striatum by immunohistochemistry. Previous confocal imaging has shown that endocytic proteins have a clustered appearance in auxilin KO primary cortical neurons (Yim et al., 2010), mainly reflective of accumulation of CCVs and clathrin cages. We stained for clathrin, and unexpectedly saw that neither the clathrin intensity nor the distribution pattern was altered in the dorsal striatum of young auxilin KO mice (Figure. 6a, b). Hsc70, the chaperone partner of auxilin, and endophilin-A1, another key endocytic protein required for uncoating also did not change significantly both in expression (Figure. 6a, d) or distribution (Figure. 6a, e) in auxilin KO mice striatum at 3 months. These observations remained true even at 9 months, except for endophilin-A1 which was significantly upregulated in auxilin KOs (Figure. 6a, c). We assessed the interaction of clathrin with Hsc70 and endophilin-A1 by their co-localization and determined it was not altered in auxilin KOs (Figure. 6a, f, g). Overall, there are no major changes in endocytic protein composition and distribution in young auxilin KO mice, in line with our synaptosomal proteomic experiments (Figure. 3e-h).

To confirm our immunohistochemistry findings, we performed EM of the dorsal striatum of 3- month-old WT and auxilin KO mice and quantitated the number of CCVs and SVs per synapse. We quantified these organelles in both asymmetric or Type I synapses which are predominantly glutamatergic (Figure. 6h), and symmetric or Type II synapses which are known to be DA or GABAergic in this brain region (Harris and Weinberg, 2012) (Figure. 6i). The number of CCVs in Type I synapses showed a moderate increase in auxilin KOs (∼10%, Supplementary Figure. 9b). Increase in CCV numbers were more pronounced in Type II synapses of auxilin KO mice (∼27%, Supplementary Figure. 9c). No notable difference between WT and auxilin KO were found in SVs number in both Type I and II synapses (Supplementary Figure. 9d, e). Cumulative effect of this was seen in CCV/SV ratio, which showed a modest but significant increase only in Type II synapses (Figure. 6k) but not in Type I synapses (Figure. 6j). It is worth noting that the distribution of CCVs and SVs was variable within the Type II synapses of auxilin KOs (Compare Figure. 6i and Supplementary Figure. 9a). Overall, these results are in consistent with our immunohistochemistry which did not show notable difference in clathrin distribution (Figure. 6a,b), and previous findings on cerebellar presynapses of auxilin KO mice (Yim et al., 2010).

The lack of a significant CCV accumulation in dorsal striatum was puzzling. To investigate whether CCVs and clathrin cages were being cleared by autophagy as suggested by several recent papers (Binotti et al., 2015; Yang et al., 2022), we examined the electron micrographs of WT and auxilin KO dorsal striatal presynapses for double membraned synaptic autophagosomes (Figure. 6l). The number of autophagosomes per synaptic site was significantly higher in auxilin Kos compared to WT (Figure 6l, m). Both Type I and II synapses showed an enhanced number of autophagosomes, though Type II synapses revealed a relatively greater number (Supplementary Figure. 8f, g). Furthermore, we find several examples of autophagosomes, in the Type II synapses, containing CCVs and SVs as their cargo (Figure. 6l, arrowheads). This unexpected finding suggests that CCVs are cleared by presynaptic macroautophagy in auxilin KO mice. These observations also support our IPA analyses of the 9-month synaptosomal proteomics data which suggested activation of macroautophagy-related pathways (Supplementary Figure. 3c).

### CCVs proteomics in auxilin KO mice suggest SV sorting defects

To understand the impact of loss of auxilin on SV sorting and composition, we performed EM and proteomic analysis of CCVs purified from brains of WT and auxilin KO mice (age: 3 months)(Blondeau et al., 2004). EM of the CCV preparations revealed that they contain both CCVs (arrows) and clathrin cages (arrowhead) but lack other organelles such as SVs (Figure. 7a) (Vargas et al., 2014). Auxilin KO mice displayed clathrin structures (CCVs + clathrin cages) which were significantly smaller in size compared to WT (Figure. 7a, b). This is in part because there was a larger proportion of clathrin cages in auxilin KOs (Figure. 7a, c), consistent with previously published findings (Yim et al., 2010).

LFQ-MS of CCVs revealed 891 proteins common to three independent experiments, 49 of which were significantly changed, with the majority being downregulated (38 down, 13 upregulated, Supplementary Table 3). Strikingly, all the proteins that exhibit decreased levels were SV transmembrane proteins (Takamori et al., 2006; Taoufiq et al., 2020), such as SNG 1 and 2, SYP, SYT 1 and 12, SV2-A and -B (Figure. 7d, e). VGLUT1 and VGLUT2, vesicular transporters for the excitatory neurotransmitter glutamate were decreased (Figure. 7d, e). Vesicular zinc transporters such as ZNT3 and TM163 were also decreased (Figure. 7d, e). By extension, this suggests that vesicular monoamine transporter-2 (VMAT2) may also be decreased, which was not detected by LFQ-MS, due to its low abundance (Taoufiq et al., 2020). IPA analysis of the CCV proteomics revealed dysregulation in the CME pathway in auxilin KO mice (Figure. 7f).

To rule out the possibility that the downregulation of certain SV transmembrane proteins seen in the auxilin KO CCV proteomics was due to the presence of clathrin cages, we compared our CCVs proteomics data with published SV proteomics data (Takamori et al., 2006). We find that the levels of synapsins, SCAMPs, syntaxins, SNAPs and several others which are categorized as SV trafficking proteins were unchanged. Endocytic proteins such as dynamins, flotilins, RAB proteins, endophilin-A1, synaptojanin-1, that are peripherally associated with the SV membrane were also unchanged in auxilin KO CCVs compared to WT (data not shown). These observations indicate that the decrease in certain SV transmembrane proteins in auxilin KO mice is likely not an artifact due to the presence of empty clathrin cages. Overall, these results suggest SV sorting defects congruent with recently proposed roles for auxilin in endocytic proof reading (Chen et al., 2019) and indicate that uncoating of CCVs in auxilin KO by GAK or alternative ways would result in SVs with an improper protein stoichiometry.

## DISCUSSION

Recent advancements in PD genetics have led to the identification of mutations in proteins that play crucial roles in SV endocytosis (Gialluisi et al., 2021; Vidyadhara et al., 2019). Mutations in clathrin uncoating proteins auxilin (*PARK19*) and synaptojanin-1 (*PARK20*) were identified to be disease-causing; while, sequence variants in GAK, and endophilin-A1 that aids in the recruitment of synaptojanin-1 to CCVs, increase risk of developing PD. Mutations in receptor-mediated endocytosis 8 (RME-8; *PARK21*), which facilitates the formation of endosome-derived SVs through clathrin uncoating, is also linked to PD (Lopert and Patel, 2014). These genetic findings strongly point to disruptions in clathrin uncoating as important for the pathogenesis of PD.

Auxilin KO mice are the first endocytic mutants to faithfully replicate the cardinal features of PD. The only other murine model with an auxilin loss-of-function mutation (R927G) displayed moderate behavior deficits but was not accompanied by DA neurodegeneration (Roosen et al., 2021). Mice with a R258Q mutation in synaptojanin-1 also displayed motor behavior deficits with no loss of DA neurons or neuroinflammation (Cao et al., 2017). In *Drosophila* loss-of-function models for the GAK homolog, *auxilin,* and RME-8 mutations, PD phenotypes are only apparent with induced α-synuclein overexpression (Song et al., 2017; Yoshida et al., 2018). Thus, endocytic mutants, besides auxilin KOs, capture only a few aspects of PD. Here, we took advantage of the fact that auxilin has a defined function in clathrin uncoating, and auxilin KOs are a robust and faithful model of PD to elucidate the underlying mechanisms. We show that auxilin deficiency leads to neurodegeneration through three distinct, but overlapping mechanisms, in nigrostriatal DA termini: 1) Toxic accumulation of cytoplasmic dopamine due to imbalance in CCV/SV ratio and defective sorting of SVs, 2) Mis-trafficking of DAT that traps the protein in axonal membrane whirls leading to defective dopamine reuptake, and 3) Synaptic autophagy overload. Collectively, these mechanisms lead to dopamine dys-homeostasis, a trigger of neurodegeneration in PD (Figure. 8).

### Accumulation of cytoplasmic dopamine

Dopamine is typically sequestered into SVs via VMAT2 to avoid autooxidation. Dopamine that accumulates in the cytoplasm is oxidized predominantly to DOPAL and subsequently catabolized to DOPAC. Thus, the elevated DOPAC levels observed in the dorsal striatum of presymptomatic auxilin KO mice, is an indirect measure of cytoplasmic dopamine accumulation (Karoum et al., 1994) and increased conversion to DOPAL, a mediator of dopamine-related toxicity in PD (Masato et al., 2019). The accumulation of DOPAC is likely due to two factors- an imbalance in the CCV/SV ratio, and SVs with improper composition. As an outcome of slowed CME (Yim et al., 2010), auxilin KO neurons need to utilize alternative endocytic pathways to maintain SV pools, which are known to be less stringent in protein sorting, leading to SVs of variable protein composition (Wu et al., 2014). Proteomic analysis of CCVs from auxilin KO brains confirmed this tenet and revealed a decrease in copy number of integral SV membrane proteins. Thus, auxilin mutations are likely to lead to fewer functional SVs available for neurotransmitter filling and release. This is supported by our neurochemical analysis which showed a decrease in extracellular dopamine metabolite 3-MT which suggest dopamine release defects (Waldmeier et al., 1981). The FSCV based computational model also predicted dopamine release defects, suggesting defective SV sequestration of dopamine in auxilin KOs. Though our CCV analysis was not sufficiently sensitive to detect VMAT2, there is a possibility of its downregulation in auxilin KO CCVs considering the decrease of two other key vesicular neurotransmitter transporters, VGLUT1 and VGLUT2.Vesicular dopamine uptake is known to be decreased in patients with PD and other synucleinopathies (Goldstein et al., 2011). VMAT2 deficient mice develop PD phenotypes, similar to auxilin KO mice, via cytosolic dopamine accumulation (Caudle et al., 2007). Previous studies have shown that DOPAL-modified α-synuclein oligomers form pores in SVs causing increased DA leakage into the cytoplasm (Plotegher et al., 2017). While SV sorting deficits probably occur in all types of synapses leading to neurotransmitter compartmentalization defects, the properties of dopamine catabolites like DOPAL are likely to render DA synapses vulnerable (Figure. 8a-c).

### Dopamine reuptake dysfunction and DAT mislocalization in membrane deformities

Extracellular dopamine in the synaptic cleft is cleared by reuptake into the presynaptic sites through plasma membrane DATs and/or enzymatic degradation to 3-MT by COMT. The relative contribution of DAT and COMT to clear extracellular dopamine varies by brain region. In dorsal striatum, reuptake by DAT plays a major role in clearing extracellular dopamine, whereas COMT has negligible role (Myöhänen et al., 2010; Yavich et al., 2007). Nigrostriatal DA presynapses depend heavily on DAT-mediated dopamine reuptake to replenish their readily releasable neurotransmitter pool. A significant delay in clearing evoked dopamine from the dorsal striatum *in vivo* and *in silico* along with presence of large DAT+ve deformities both in fixed tissue and *ex vivo* clearly indicate DAT dysfunction in auxilin KO mice. This appears to be a defining feature of DA neurodegenerative phenotypes in auxilin KOs. Dopamine reuptake dysfunction for an extended period may lead to striatal dopamine loss as seen in DAT KOs (Jones et al., 1998), and exacerbate PD pathology in auxilin KO mice.

Live slice imaging of rhodamine-tagged dichloropane which binds to plasma membrane DAT, as well as the DAT-immunogold labelling in the dorsal striatum confirmed DAT-rich axonal membrane deformities in auxilin KOs. Similar DAT positive membrane whirls have been described for synaptojanin-1 PD mutants (Cao et al., 2017), and may be a common feature of endocytic PD mutants. Other evidence of axonal damage comes from our proteomic findings as well, where neurofilament proteins that maintain axonal integrity were altered, including an increase in NF-L, a marker of neuroaxonal damage. In DA presynapses, DAT localization to the plasma membrane is dynamically regulated by endocytic trafficking and recycling. It remains to be determined whether auxilin and synaptojanin-1 participate in endocytic recycling of DAT in the DA neurons and will be explored in the future. However, our observations suggest that the dopamine reuptake decrement seen in auxilin KOs occur principally due to DA axonal membrane deformities which trap DAT (Figure. 8d).

### Synaptic autophagy overload

Owing to the higher turnover of synaptic proteins, vesicles, and mitochondria in presynaptic sites, autophagosome biogenesis occurs at a higher rate in the distal axons than in the soma (Maday and Holzbaur, 2014). Due to limited lysosomal activity, synaptic termini depend on retrograde microtubule-based axonal transport of autophagosomes towards the lysosome-rich soma for degradation. Tonic activity of DA neurons is likely to keep basal autophagy rates high, and the requirement to transport autophagosomes long distances via extensive arborization make DA axons vulnerable to additional autophagic burden. Ultrastructural evaluation of dorsal striatum of auxilin KO mice revealed an increase in synaptic autophagosomes, which was more pronounced in Type II synapses. We also observed autophagosomes containing CCVs and SVs in auxilin KO synapses. We find evidence for increased mTOR signaling and activation of autophagic pathways in the synaptosomal proteomic data at symptomatic age, supporting elevated synaptic autophagy in auxilin KO mice. Rapamycin-induced enhancement of autophagy in DA presynapses of mice striatal slices have been shown to sequester SVs and decrease evoked dopamine release (Hernandez et al., 2012). A similar event in auxilin KO synapses might exacerbate cytosolic dopamine accumulation (Figure. 8a). Enhanced synaptic autophagy to clear missorted SVs and CCVs, as well as the products of toxic dopamine-oxidation could overload DA projections with autophagic vacuoles. Autophagosome accumulation, impaired retrograde transport as well as abnormal axonal deformities in DA axons have been previously noted in neurons from patients with PD and Alzheimer’s disease (Hill and Colón-Ramos, 2020; Kouroupi et al., 2017; Nixon et al., 2005). EM revealed some of the autophagic vacuoles near the whirl-like axonal deformity in auxilin KO striatum. Though we presently do not understand the relationship between these two structures, DA axonal deformities observed in auxilin KOs may be a result of autophagic overload in DA termini.

In conclusion, our findings indicate that pathology of PD mediated by auxilin deficiency begin with a disruption of CME, which leads to fewer functional SVs for neurotransmitter filling. While these deficits occur at all synapses, it appears to have a particularly detrimental effect at nigrostriatal DA synapses due to the toxicity of cytosolic dopamine, and DAT reuptake alterations. Thus, investigating auxilin loss-of-function has also advanced our understanding of the mechanisms for DA vulnerability of PD.

## STAR METHODS

### Mice

Auxilin KO mice have been previously described (Yim et al., 2010) and were bred to C57BL6/J mice to make them congenic. Auxilin homozygous KOs were compared to WT C57BL6/J from Jackson Laboratories, Maine. We have an IACUC approved protocol to maintain these mice.

### Motor behavior evaluation

WT and auxilin KO cohorts were examined longitudinally at 3, 6, 9, 12, and 15 months of age (n=12-16 mice/genotype, sex-balanced) in motor behavioral assays. For evaluation of overall locomotory capabilities, mice were allowed to explore an open field arena, which was videotaped to assess the distance travelled in 5 mins using Noldus Ethovision CT software. The number of fecal pellets excreted during open field behavior test was evaluated as a measure of anxiety. The balance beam test was used to assess motor coordination by evaluating the ability to walk straight on a narrow beam from a brightly lit end towards a dark and safe box. Latency to traverse the beam and the number of times a mouse could perform this behavior in a minute were evaluated. The grip strength of the forelimbs and all the limbs was assessed by measuring the maximum force (g) exerted by the mouse in grasping specially designed pull bar assemblies attached to a grip strength meter (Columbus Instruments, Ohio, USA) in tension mode. A four-lane Rotarod was used to assess motor coordination and balance (Columbus Instruments, Ohio, USA). Mice were made to run for 300 secs on the rotating spindle of the Rotarod, which was accelerating from 4 to 40 rpm. Each mouse was subjected to three trials with a 30 min inter-trial recovery period. The average of the latency to fall and the rpm in these trials was used as a measure of motor performance. The procedure was repeated for four consecutive days in both WT and auxilin KO mice. We did not see a significant sex-based differences in all the behavior assays in auxilin KO mice and data from both sexes was collated.

### Immunohistochemistry

WT and auxilin KO mice at 3 and 9 months of age (n=5-6/group; sex-balanced) were anaesthetized using isoflurane inhalation and perfused intracardially with 0.9 % heparinized saline followed by chilled 4 % paraformaldehyde (PFA) in 0.1 M phosphate buffer (PB). The brains were post-fixed in the same buffer for 48 hours and cryoprotected in increasing grades of buffered sucrose (15 and 30 %, prepared in 0.1 M PB), at 4 °C, and stored at −80 °C until sectioning. Serial sections of the brains (30 μm thick) were performed coronally using a cryostat (Leica CM1850, Germany), collected on gelatinized slides, and stored at -20 °C. Every sixth nigral section was subjected to immunoperoxidase staining and every 10^th^ striatal section was used for immunofluorescence staining as per our earlier protocol(Vidyadhara et al., 2017). Briefly, for immunoperoxidase staining, endogenous peroxidase quenching was performed using 0.1 % H2O2 in 70 % methanol (30 mins), followed by blocking using 3 % bovine serum albumin (BSA) (2 hours) at room temperature (RT). Sections were then incubated at 4 °C with TH primary antibody (1:500, overnight) followed by biotin-conjugated secondary antibody at RT (1:200; 3-4 hours, Vector Laboratories, PK-6101). Tertiary labeling was performed with the avidin-biotin complex solution at RT (1:100; 3-4 hours, Vector Laboratories, PK-6101). Staining was visualized using 3,3’-diaminobenzidine (Fluka, 32750) as a chromogen in a solution of 0.1 M acetate imidazole buffer (pH 7.4) and H2O2 (0.1 %). For immunofluorescence staining, sections were incubated in 0.5 % Triton-X-100 (15 mins), followed by incubation in 0.3 M glycine (20 mins). Blocking was performed using 3% goat serum, followed by overnight incubation (4° C) in primary antibodies. Sections were then incubated in Alexa-conjugated secondaries (Thermo Fisher Scientific, USA) for 3-4 hours, followed by coverslip mounting using an antifade mounting medium with (H-1000, Vectashield) or without DAPI (H-1200, Vectashield). Coverslips were sealed using nail polish. 1X PBS with 0.1 % Triton-X-100 was used as both washing and dilution buffer for both immunoperoxidase and immunofluorescence staining, except for pSer129-α-Syn where 1X Tris buffer saline was used. Below is the list of antibodies used and their dilutions.

| Antibody | Dilution | Manufacturer | RRID |
| --- | --- | --- | --- |
| Rabbit Anti-TH | 1:500 | Millipore (AB152) | RRID: AB_390204 |
| Mouse Anti-TH | 1:500 | Synaptic Systems (213211) | RRID: AB_2636901 |
| Rabbit Anti-Iba1 | 1:300 | Wako Chemicals (019-19741) | RRID: AB_839504 |
| Guinea Pig Anti-GFAP | 1:400 | Synaptic Systems (173004) | RRID: AB_10641162 |
| Guinea Pig Anti-DAT | 1:300 | Synaptic Systems (284005) | RRID: AB_2620019 |
| Rabbit Anti-α-synuclein (phospho S129) | 1:800 | Abcam (ab51253) | RRID: AB_869973 |
| Mouse Anti-Clathrin light chain | 1:200 | Synaptic Systems (113011) | RRID: AB_887706 |
| Rabbit Anti-Synaptogyrin 3 | 1:200 | Synaptic Systems (103303) | RRID: AB_2619753 |
| Rabbit Anti-VGAT | 1:500 | Synaptic Systems (131002) | RRID: AB_887871 |
| Guinea Pig Anti-VGLUT2 | 1:500 | Synaptic Systems (135418) | RRID: AB_2864786 |
| Rat Anti-Hsc70 | 1:100 | Enzo (ADI-SPA-815-D) | RRID: AB_2039279 |
| Rabbit Anti-Endophilin-A1 | 1:200 | Synaptic Systems (159002) | RRID: AB_887757 |

### Unbiased stereology

The SNpc was delineated on every 6^th^ TH+ve midbrain section(Fu et al., 2012) using a 10X objective of a brightfield microscope equipped with StereoInvestigator (Software Version 8.1, Micro-brightfield Inc., Colchester, USA). The stereological quantification of TH+ve DA neurons was performed using the optical fractionator probe of the StereoInvestigator(Vidyadhara et al., 2017). The neurons were counted using 40X objective, with a regular grid interval of 22,500 μm² (x=150 μm, y=150 μm) and a counting frame size of 3600 μm² (x=60 μm, y=60 μm). The mounted thickness was identified to be around 22.5 μm, which was also determined at every fifth counting site. A guard zone of 3.5 μm was implied on either side, thus providing 15 μm of z-dimension to the optical dissector. The quantification began at the first anterior appearance of TH+ve neurons in SNpc to the caudal most part in each hemisphere(Fu et al., 2012) separately, which was later summed to derive total numbers.

### Confocal microscopy and image analysis

Fluorescent images were acquired using a laser scanning confocal microscope (LSM 800, Zeiss) with a 20X or 40X objective for quantitation and 63X for representation using an appropriate Z-depth. All the images were blinded for genotype and age before subjecting to analysis using FIJI software from National Institute of Health (NIH). After performing sum intensity projection, the expression intensity was measured on an 8-bit image as the mean gray value on a scale of 0–255, where ‘0’ refers to minimum fluorescence and ‘255’ refers to maximum fluorescence. For counting Iba1+ve microglial cells, images were thresholded using the ‘otsu’ algorithm and the cells larger than 75-pixel units for a given image were counted using the ‘analyze particles” function. GFAP+ve astroglial cells were counted manually using the ‘cell counter’ function. For counting DAT+ve structures, images were thresholded using ‘triangle’ algorithm, followed by ‘analyse particles’ function. All the structures of size 5 µm and above and the circulating between 0.3 to 1 were counted. SNpc, VTA and SNpr were demarcated as per Fu et al., 2012, colabeling with TH-immunostaining (Fu et al., 2012). Dorsal and ventral striata were demarcated as per Paxinos and Franklin, 2008 (Keith Franklin, 2008).

### Proteomic analysis

Whole brain, synaptosomes and CCV samples were prepared from 3-month-old WT and auxilin KO mice. Brains from 3-month-old WT and auxilin KO mice (n=3/genotype) were homogenized in homogenization buffer (detergent-free 320 mM sucrose in 10 mM HEPES, pH 7.4 with protease and phosphatase inhibitors cocktail). Part of the homogenate was snap-frozen for whole brain proteomics. Rest of the homogenate was used to prepare synaptosomes as described previously (De Camilli et al., 1983). Synaptosomes integrity was confirmed by EM before performing LFQ-MS. For CCVs sample preparation, brains from 14 pairs of WT and auxilin KO mice were pooled to obtain a single CCV fraction (Blondeau et al., 2004) as we have published previously (Vargas et al., 2014). Three independent purifications were performed and the resulting CCVs fractions were subjected to LFQ-MS. The purity of CCVs was confirmed by EM (Figure. 7a).

LFQ-MS was performed at Yale Mass Spectrometry & Proteomics Resource of the W.M. Keck Foundation Biotechnology Resource Laboratory. Samples were analyzed in technical triplicates. The raw mass spectrometery data will be publicly available upon publication in the PRIDE depository. The data was normalized to internal controls and total spectral counts. Proteins with two or more unique peptide counts were listed using UniProt nomenclature and included for further analysis. A 1.5-fold change and a p-value difference of <0.05 between WT and auxilin KO are considered as significant. Heat maps for significantly changed proteins were produced using Qlucore Omics Explorer. IPA (Qiagen) was used to determine the most significantly affected canonical pathways and their overlap.

### High-performance liquid chromatography (HPLC)

Sex balanced, auxilin KO mice at 3, 6, 9, 12, and 15 months of age with appropriate controls (n=8-12/genotype) were anesthetized using isoflurane inhalation. Mice were then sacrificed by cervical dislocation, and the brains were quickly removed and dissected for dorsal striatum. These samples were subjected to HPLC at Vanderbilt Neurochemistry Core Laboratory, Vanderbilt University. We did not notice a significant sex-based difference in HPLC results in auxilin KO mice.

### Surgery and *in vivo* Fast Scanning Cyclic Voltammetry (FSCV)

Surgeries and electrochemical recordings were conducted like our published procedure(Somayaji et al., 2020). Briefly, mice were anesthetized with isoflurane (SomnoSuite Small Animal Anesthesia System, Kent Scientific; induction 2.5%, maintenance 0.8–1.4% in O^2^, 0.35 l/min) and head-fixed on a stereotaxic frame (Kopf Instruments, Tujunga, CA). Puralube vet ointment was applied on the eye to prevent cornea from drying out. Stereotactic drill (0.8 mm) was used to preform craniotomy (unilateral, right) to target the midbrain and dorsal striatum with the following coordinates(Keith Franklin, 2008) (values are in mm from Bregma); midbrain: anteroposterior = -2.9, mediolateral=+1.0, dorsoventral=+4; Dorsal Striatum: anteroposterior = +1.2, mediolateral = +1.3, dorsoventral = +3.1. An Ag/AgCl reference electrode via a saline bridge was placed under the skin. For electrical stimulations, a 22G bipolar stimulating electrode (P1 Technologies, VA, USA) was lowered to target ventral midbrain (between 4-4.5mm). The exact depth was adjusted for maximal dopamine release. For recording the evoked dopamine release, a custom-built carbon fiber electrode (5 μm diameter, cut to ∼150 μm length, Hexcel Corporation, CT, USA) was lowered to reach dorsal striatum. Dil-coated carbon-fiber electrodes were used to identify the electrode position in the dorsal striatum and the electrode track in the brain tissue identified the position of the stimulation electrode (Supplementary Figure. 3c). The evoked dopamine release was measured using constant current (400μA), delivered using an Iso-Flex stimulus isolator triggered by a Master-9 pulse generator (AMPI, Jerusalem, Israel). A single burst stimulation consisted of 30 pulses at 50Hz (0.6s). Electrodes were calibrated using known concentration of dopamine in ACSF. Custom-written procedure in IGOR Pro was used for the data acquisition and analysis.

### Computational model of dopamine reuptake and release

The model is comprised of a system of ordinary differential equations (ODEs), where equations (1) and (3) form a two-compartment model to simulate the release of dopamine (DA) from synaptic vesicles into the dorsal striatum and the diffusion of DA towards the carbon-fiber electrode through a “dead space” (i.e. an area of damaged tissue around the carbon fiber electrode) (Benoit-Marand et al., 2007):

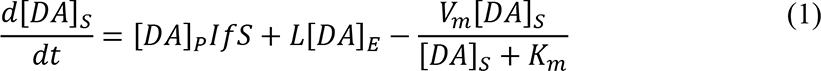

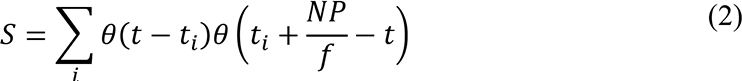

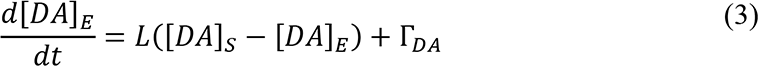

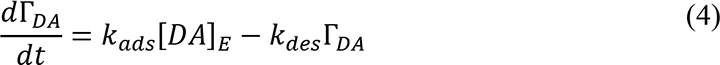

The first compartment [*DA*]_*s*_ sums the concentration of DA released into the striatum, the concentration of DA that returns to the carbon-fiber electrode due to its reflective surface(Schmitz et al., 2001), and the concentration of DA removed through reuptake by DAT. [*DA*]_*p*_ is the concentration of DA release per electrical stimulus pulse, while *I* and *f* are the stimulus current and stimulus frequency of the experimental protocol. The electrical pulse trains are modeled using the stimulation pattern *S*, where θ is the Heaviside theta function, *t*_*i*_ is the start time of the stimulus, and *NP* is the number of electrical pulses. DAT uptake is modeled using first-order Michaelis-Menten kinetics (Michaelis et al., 2011; Wightman and Zimmerman, 1990), with *v*_*m*_ and *k*_*m*_ as the maximal velocity and affinity constant of DA. The second compartment [*DA*]_*E*_ computes the difference between the DA that arrives from the striatum and the DA that bounces off the electrode, with a loss factor *L* < 1 used to account for the diffusion through the dead space. Equation (4) is used to model the electrochemical adsorption that occurs with carbon-fiber electrodes(Bath et al., 2000) and is included in the calculation of [*DA*]_*g*_, and *k*_*ads*_ and *k*_*des*_ are the adsorption and desorption kinetic rate constants.

### *Ex vivo* dichloropane-DAT imaging and quantitation

Dichloropane, a DAT ligand, was conjugated with rhodamine red-X as described by Fiala et al., 2020 (Fiala et al., 2020) to obtain dichloropane–rhodamine red-X probe (dichloropane probe). Mice (n=5/genotype) were sacrificed by cervical dislocation under isoflurane anesthesia and the brains were quickly dissected. Coronal slices (300 µm) of striatum were cut (VT1200S, Leica) in ice-cold carbogenated solution containing: 100 mM choline chloride, 25 mM NaHCO3, 1.25 mM NaH2PO4, 2.5 mM KCl, 7 mM MgCl2, 0.5 mM CaCl2, 15 mM glucose, 11.6 mM sodium ascorbate, and 3.1 mM sodium pyruvate. Striatal slices were incubated at 37°C (30 mins) in carbogenated ACSF containing: 127 mM NaCl, 25 mM NaHCO3, 1.25 mM NaH2PO4, 2.5 mM KCl, 1 mM MgCl2, 2 mM CaCl2 and 15 mM glucose. Slices were then warmed to room temperature in carbogenated ACSF and incubated in dichloropane probe (100 nM) for 45 mins at 37°C. Following this, slices were washed once in carbogenated ACSF and the dorsal striata maintained in ACSF were imaged under 25X water immersion objective at 561 nm excitation using a confocal microscope (LSM 900, Zeiss). Images were analyzed blind to the genotype for the presence of large DAT+ve structures and the number of DAT+ve puncta using FIJI software.

### Electron microscopy

3-month-old mice brains (n=2-3/genotype) were fixed by intracardial perfusion using 2% PFA and 2% glutaraldehyde prepared in 0.1M PB, followed by overnight immersion in 0.1 M cacodylate buffer with 2.5 % of glutaraldehyde and 2 % PFA. For DAT- immunogold labeling (15 nm gold particles), we used 3% PFA prepared in 1X PBS for intracardial perfusion, and 2% PFA and 0.15% glutaraldehyde prepared in 1X PBS for immersion fixation. Dorsal striatum was dissected, further processed at the Yale Center for Cellular and Molecular Imaging, Electron Microscopy Facility. EM imaging was performed using FEI Tecnai G2 Spirit BioTwin Electron Microscope. Images were analyzed blinded to the genotype using FIJI software for synaptic autophagosomes, both in symmetric and asymmetric synapses. Similarly, synaptic CCVs and SVs per synapse were also counted, along with examining the images for axonal whirls and early autophagic vacuoles.

For EM of purified CCVs and clathrin cages, buffer containing CCVs was pipetted onto a parafilm containing glutaraldehyde and uranyl acetate to make a 18% glutaraldehyde and 73% uranyl acetate solution in 1X PBS. EM grids were floated on top of pipetted droplets and then dried for imaging, using Philips 301 Electron Microscope. Diameter of the CCVs and empty clathrin cages, as well as their numbers were measured using iTEM software (ResAlta Research Technologies, USA).

### Statistics

For behavioral studies, two-way repeated measure ANOVA followed by Sidak’s multiple comparison test was used. For all other experiments, Student’s t-test with Welch’s correction was used. Values are expressed as mean ± standard error of the mean (SEM) and *p* value of 0.05 or less was considered statistically significant. Student’s t-test was also used to check if there are sex-based differences in the experimental results within auxilin KO mice, which was not significant.

## ACKNOWLEDGMENTS

This work was supported by Parkinson’s Foundation Research Center of Excellence (PF-RCE-1946), Nina Compagnon Hirsheld Parkinson’s Disease Research Fund to SSC, and Michael J. Fox Foundation Target Advancement Program grant (MJFF-020160) to SSC and VDJ. SSC and DS are funded by ASAP and are members of the ASAP CRN. We thank Pietro De Camilli for his input and valuable suggestions. We acknowledge John Lee’s help in initial behavioral experiments. We thank Sofia M. Tieze and Phil Coish for reading the manuscript. We thank the Strittmatter lab for providing access to the open field behavior setup, and the Rakic Lab for the StereoInvestigator facility.

## AUTHOR CONTRIBUTIONS

VDJ, DLS and SSC conceptualized the study. VDJ performed all behavior and histochemical experiments, imaging, quantitation and proteomic analyses. MS performed *in vivo* FSCV experiments. NW performed analyses of CCV proteomics and EM images. BY performed mice genotyping. HZ prepared CCVs. SN performed computational analysis. JR helped in immunofluorescence image analysis and illustrations. JG prepared *ex vivo* brain slices. TLL performed LFQ-MS. DS and LG provided reagents and founder mice colonies. VDJ and SSC wrote the manuscript. All authors have read and provided inputs to the manuscript.

## ABBREVIATIONS

3-MT: 3-Methoxytyramine
5-HIAA: 5-hydroxyindoleacetic acid ACSF: Artificial cerebrospinal fluid ALDH7A1: Aldehyde dehydrogenase 7A1 BSA: Bovine serum albumin
CCVs: Clathrin coated vesicles
CME: Clathrin mediated endocytosis
COMT: Catechol-o-methyltransferase
DA: Dopaminergic
DAT: Dopamine transporter
DOPAC: 3,4-dihydroxyphenylacetic acid
EM: Electron Microscopy
FSCV: Fast scan cyclic voltammetry
GAK: Cyclin G-associated kinase
GFAP: Glial fibrillary acidic protein
HPLC: High performance liquid chromatography
HVA: Homovanillic acid
Iba1: Ionized calcium-binding adapter molecule 1
IPA: Ingenuity Pathway Analysis
KO: Knockout
RME-8: Receptor-mediated endocytosis 8
SNG: Synaptogyrin
SNpc: Substantia nigra pars compacta
SEM: Standard error of the mean
SV: Synaptic vesicle
SV2: Synaptic vesicle glycoprotein 2
SYP: Synaptophysin
Syn: Synuclein
SYT: Synaptotagmin I
TH: Tyrosine hydroxylase
VGAT: Vesicular GABA transporter
VGLUT: Vesicular glutamate transporters
VMAT2: Vesicular monoamine transporter-2
VTA: Ventral tegmental area
WT: Wildtype

---

**Supplementary Figure. 1:**
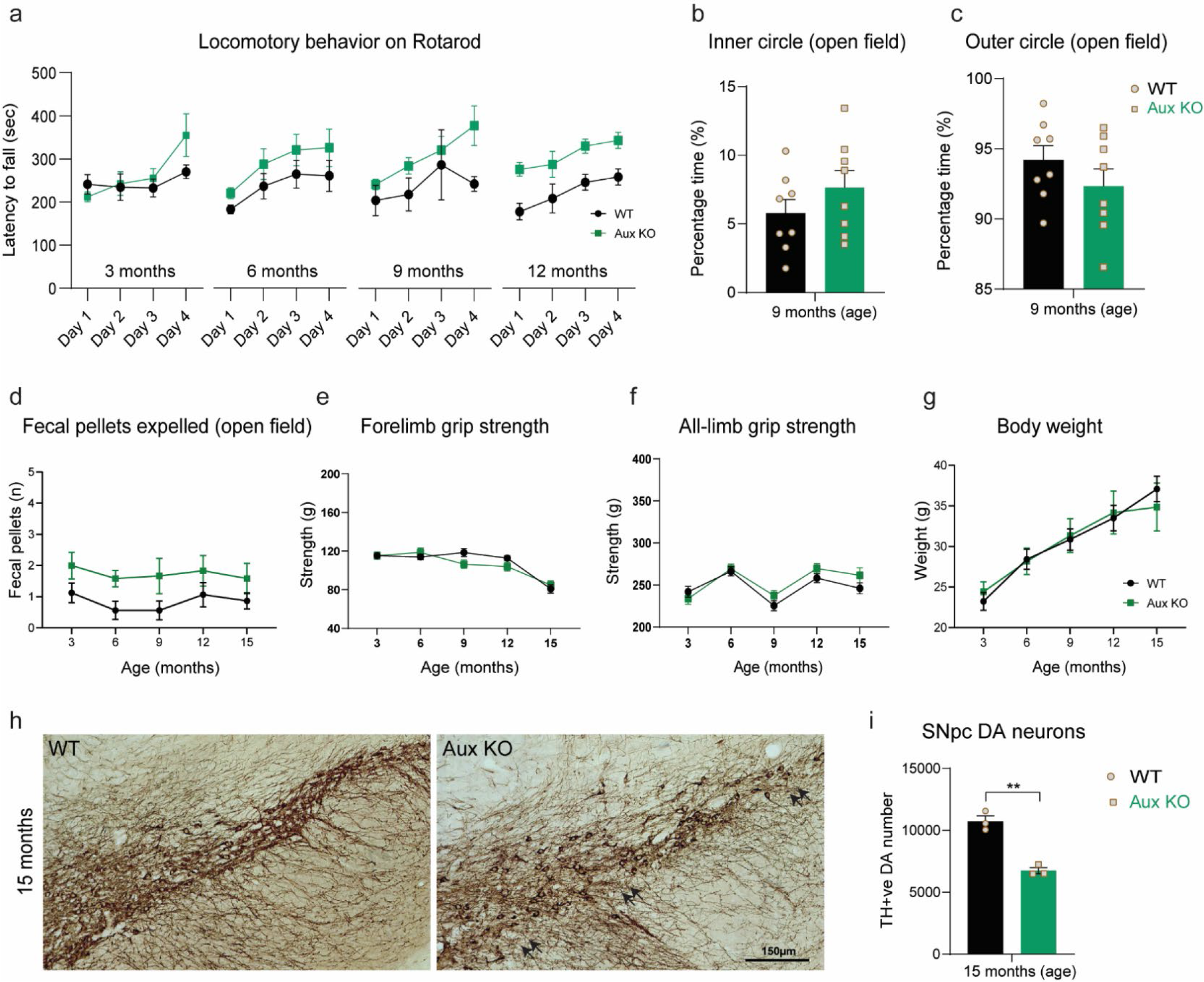
Auxilin KO mice show selective behavioral deficits. **a.** Motor coordination on Rotarod measured as latency to fall. Aux KOs did not show significant alteration in this behavioral test when compared to WT. **b.** Percentage time spent in inner circle of the open field. This is a measure of anxiety and was not different from WT at 9 months in Aux KOs, even though motor deficits were apparent. **c.** Percentage time spent in outer circle of the open field. **d.** Number of fecal pellets expelled during open field behavior, which did not change significantly between WT and Aux KOs across age. **e.** Forelimb grip strength measured as a function of age in WT and Aux KOs. **f.** All-limb grip strength was also not affected. **g.** Body weight of WT and Aux KOs as a function of age. **h.** Representative images of TH+ve DA neurons in SNpc of WT and Aux KO mice at 15 months of age. Note a loss of DA neurons in the SNpc of Aux KO mice (arrows). Scale bar: 150 μm. **i.** Stereological counting of SNpc DA neurons, which revealed a significant loss of DA neurons in Aux KO mice at 15 months of age. Statistics: For age-related behavior, two-way repeated measure ANOVA followed by Sidak’s multiple comparison test was used. For others, Student’s t-test with Welch’s correction was used. *\*\*p<0.01*.

**Supplementary Figure. 2:**
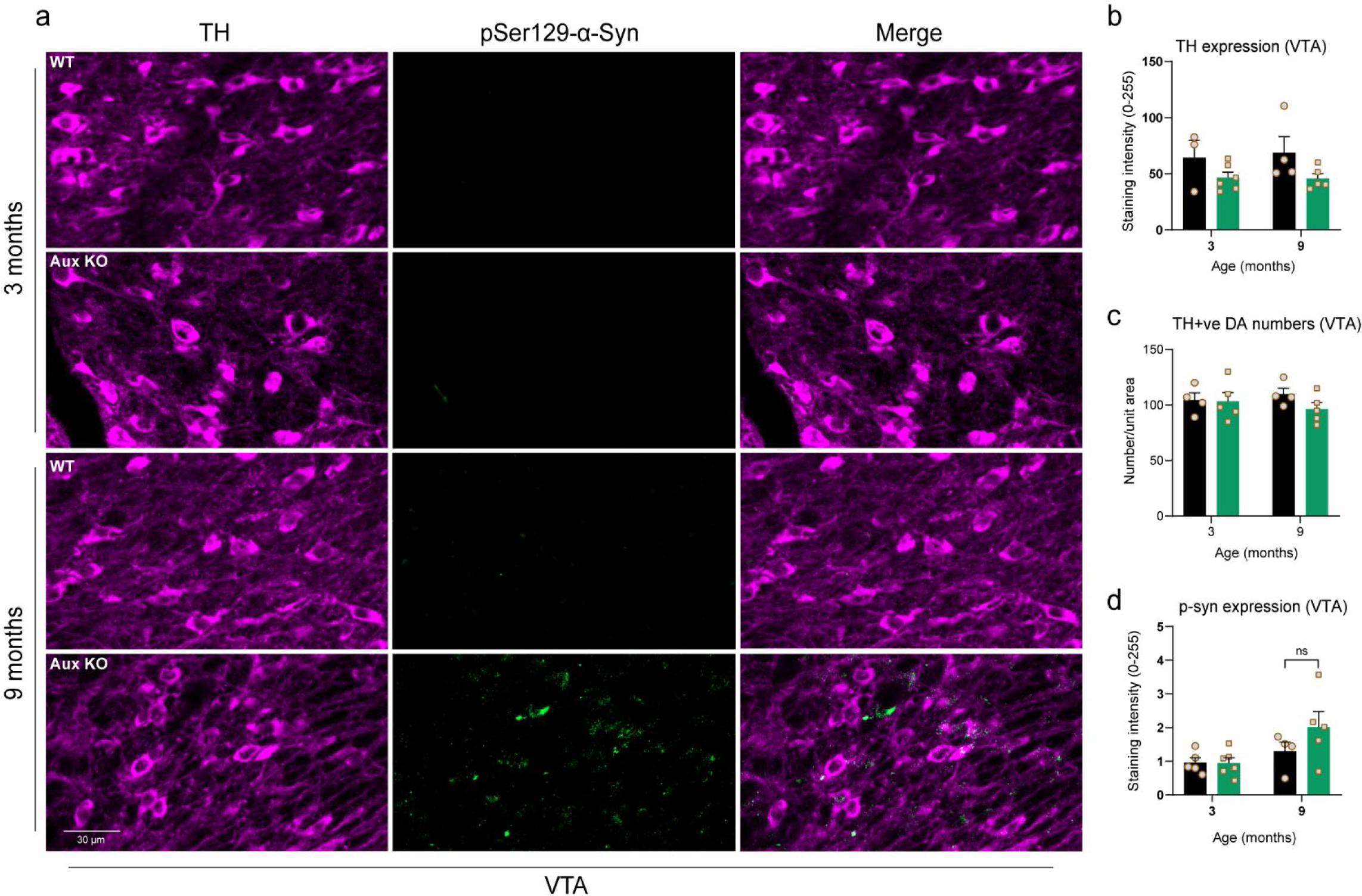
VTA was relatively preserved in auxilin KOs. **a.** Representative images VTA immunostained for α-synuclein aggregation marker pSer129-α-synuclein (green) co-labelled with DA marker TH, at 3 and 9 months in WT and Aux KOs. Scale bar: 30 μm. **b.** TH expression in VTA, which did not change with age in Aux KOs. **c.** Number of TH+ve DA neurons in VTA, which was not altered in Aux KO mice. **d.** p-Ser 129-α-synuclein expression in VTA, which showed a trend of higher expression at 9 months but did not reach significance in Aux KOs. Statistics: Student’s t-test with Welch’s correction.

**Supplementary Figure. 3:**
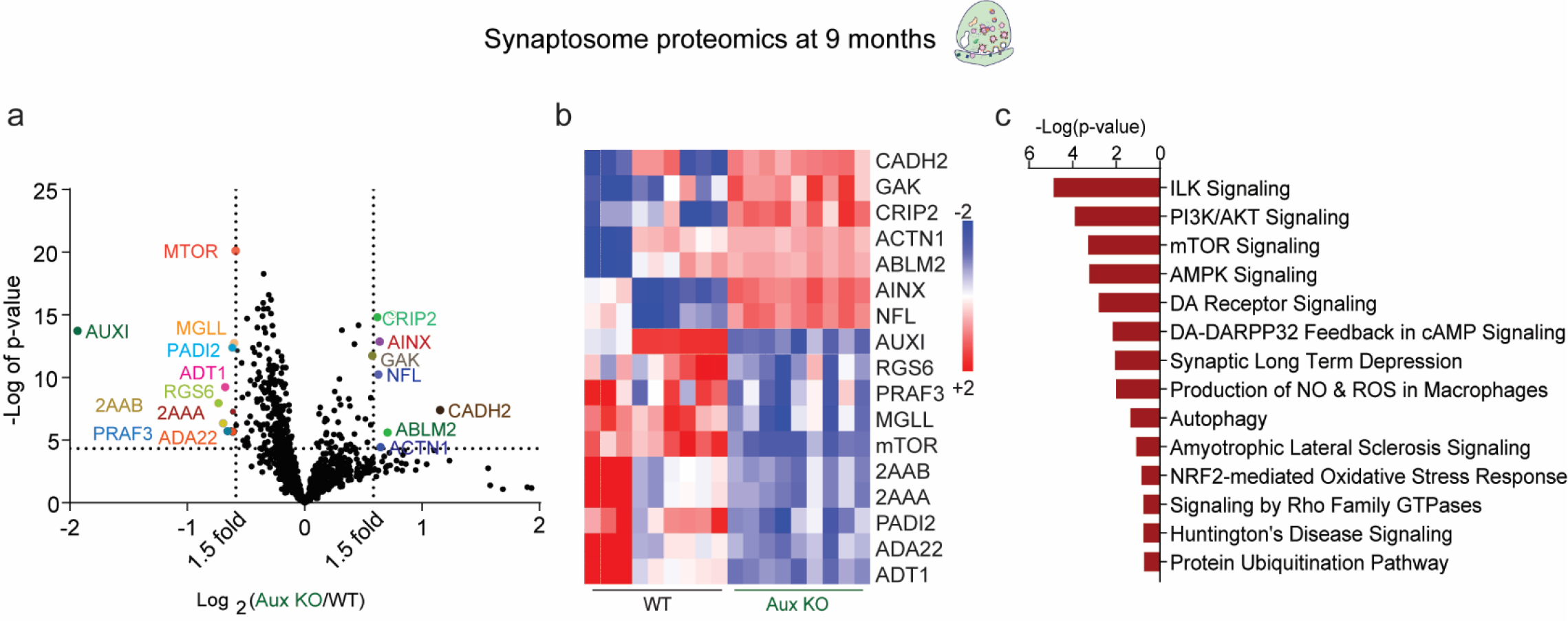
Synaptosome proteomics at 9 months revealed several PD-linked proteins and pathways to be altered. **a.** Volcano plot of synaptosome proteome of 9-month-old Aux KOs compared to WT (n=3 mice/genotype). Proteins that were altered greater than 1.5-fold (vertical dotted lines) with a p-value of 0.05 (Student’s t-test) or lesser (horizontal dotted line) were considered as significantly changed. Among 17 proteins that significantly changed, 10 were decreased (left) and 7 were increased (right). **b.** Heat map of significantly changed proteins in synaptosomes of Aux KOs in comparison to WT depicted for each technical replicate (3 technical replicates/mouse). Red indicates an increase (+2) and blue indicates decreased levels (-2). **c.** Pathways that are significantly (p<0.05) affected in whole brain synaptosomes of Aux KOs as determined by IPA.

**Supplementary Figure. 4:**
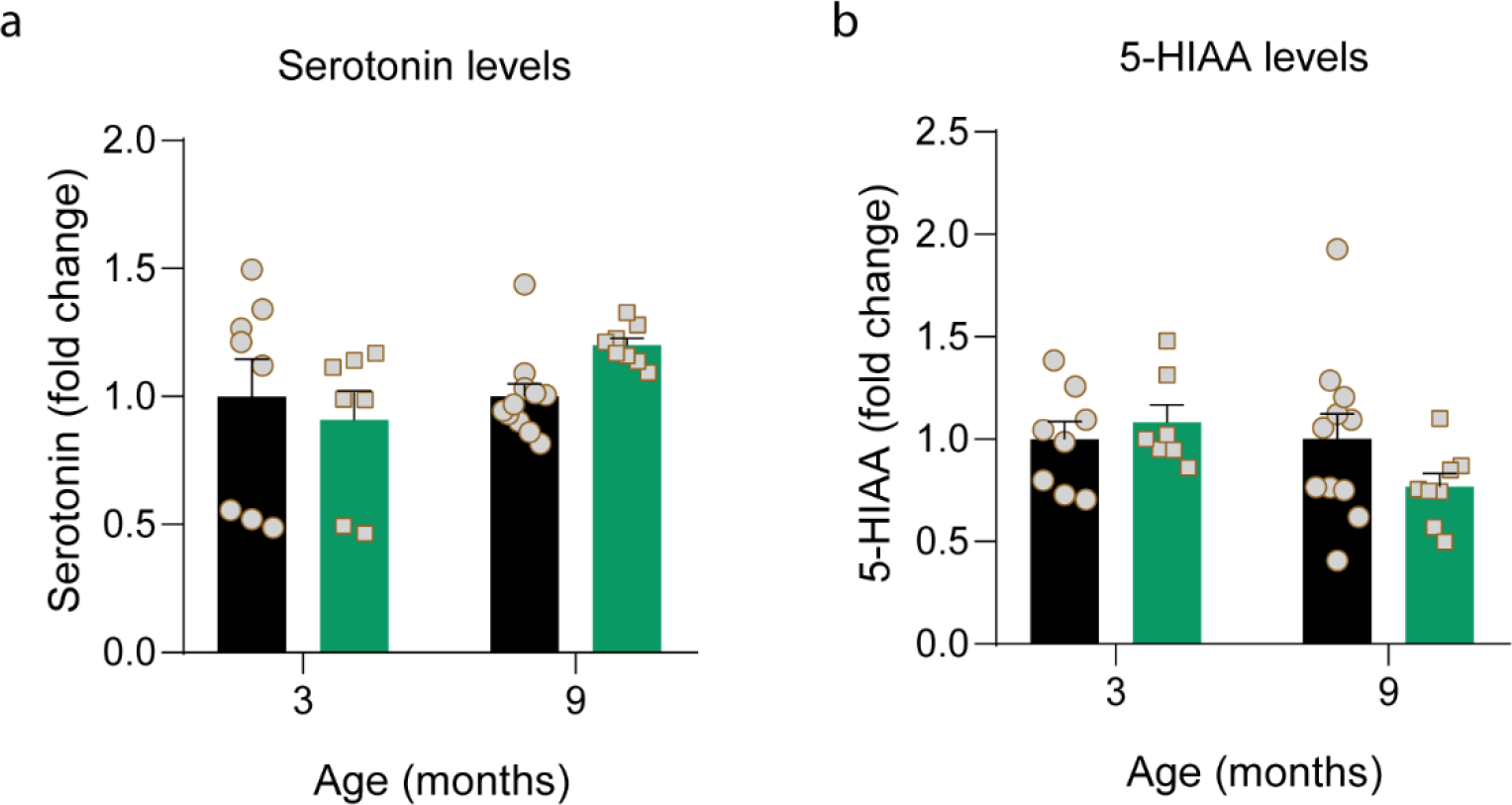
Serotonin and its metabolites were unaltered in auxilin KO brains. **a.** Serotonin levels in the dorsal striatum of WT and Aux KOs at 3 and 9 months, as measured by HPLC, which was not altered in Aux KO mice. **b.** Levels of 5-HIAA, a serotonin metabolite, was also unaltered. Statistics: Student’s t-test with Welch’s correction.

**Supplementary Figure. 5:**
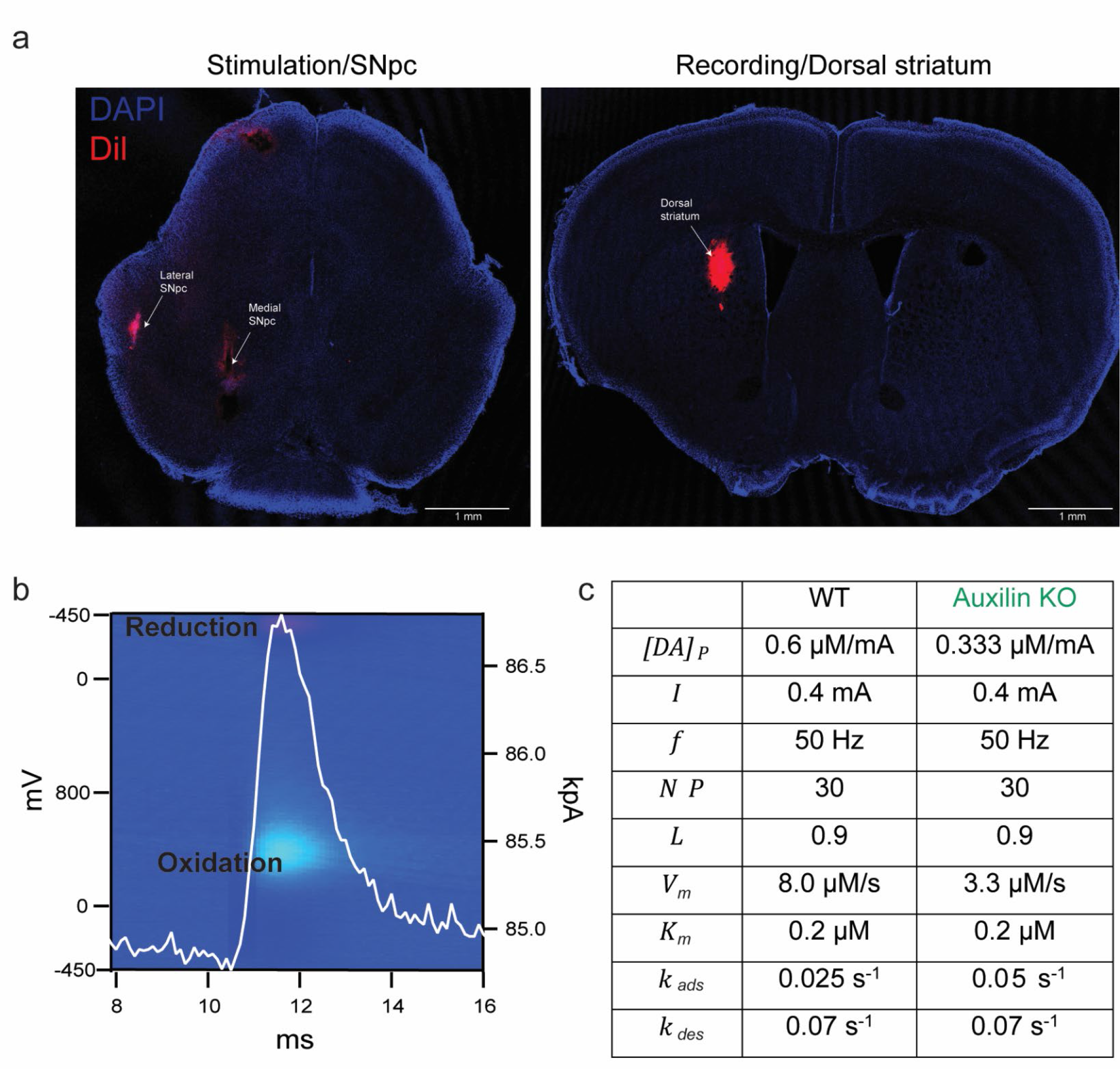
FSCV recording. **a.** Representative images of coronal mouse brain sections showing the location of the bipolar stimulating electrode in the SNpc and the FSCV recording electrode in the dorsal striatum (STR), as marked by DiI staining (DiI: red, DAPI: blue). Scale bar: 1mm **b.** The 3-dimensional pseudocolor plot showing oxidation (cyan) and reduction (red) of dopamine. **c.** Best fit parameters of the dopamine computational model to fit FSCV recordings.

**Supplementary Figure. 6:**
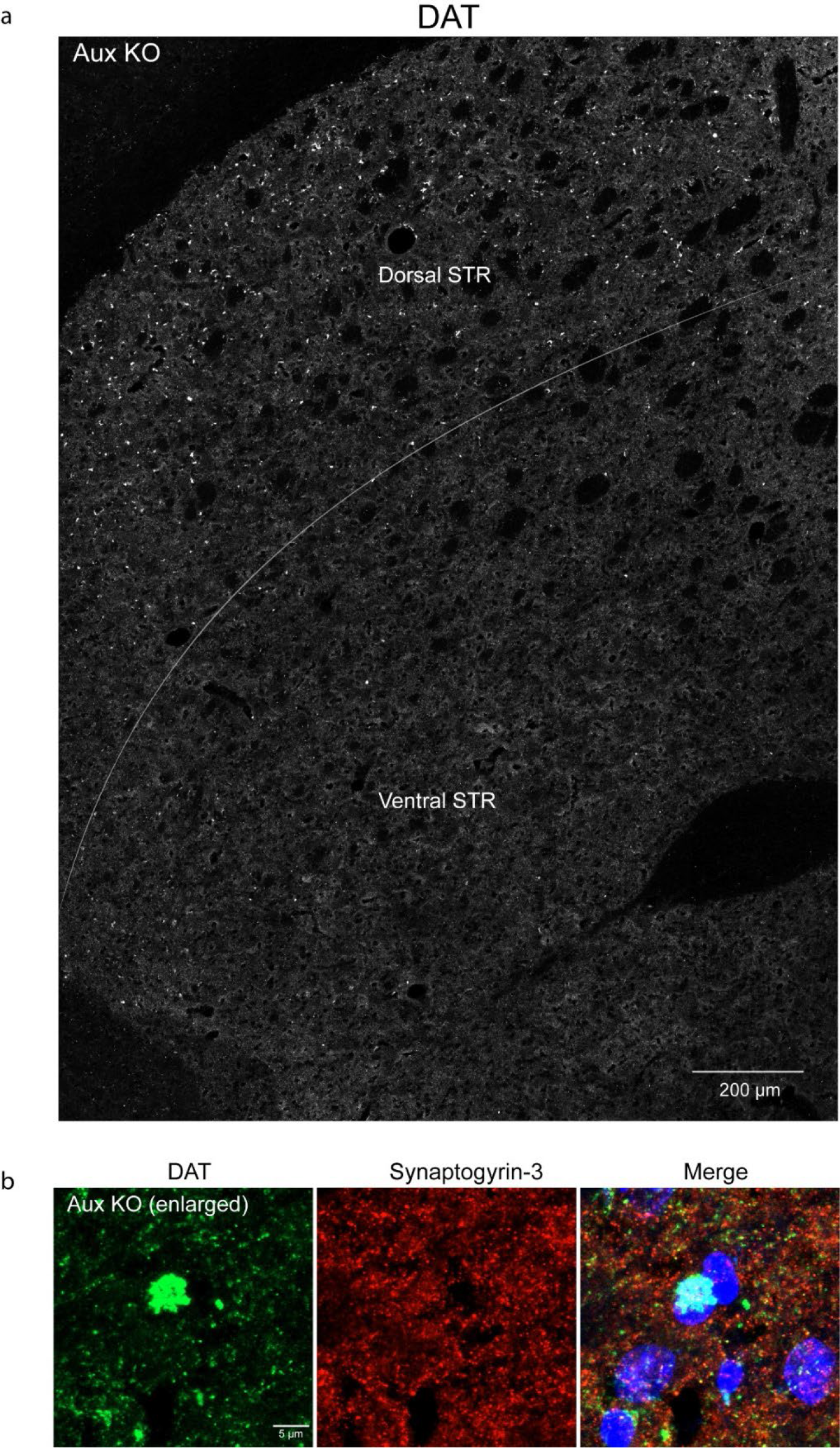
Large DAT+ structures in the dorsal striatum of auxilin KOs. **a.** Representative grayscale image of striatum of Aux KOs immunostained for DAT, showing large DAT+ structures are enriched in the dorsolateral striatum (STR), but not in the ventral STR. Scale bar: 200 μm. **b**. Enlarged image of DAT+ve structures (green) in the dorsal striatum, co- immunostained with synapogyrin-3 (red). Scale bar: 5 μm.

**Supplementary Figure. 7:**
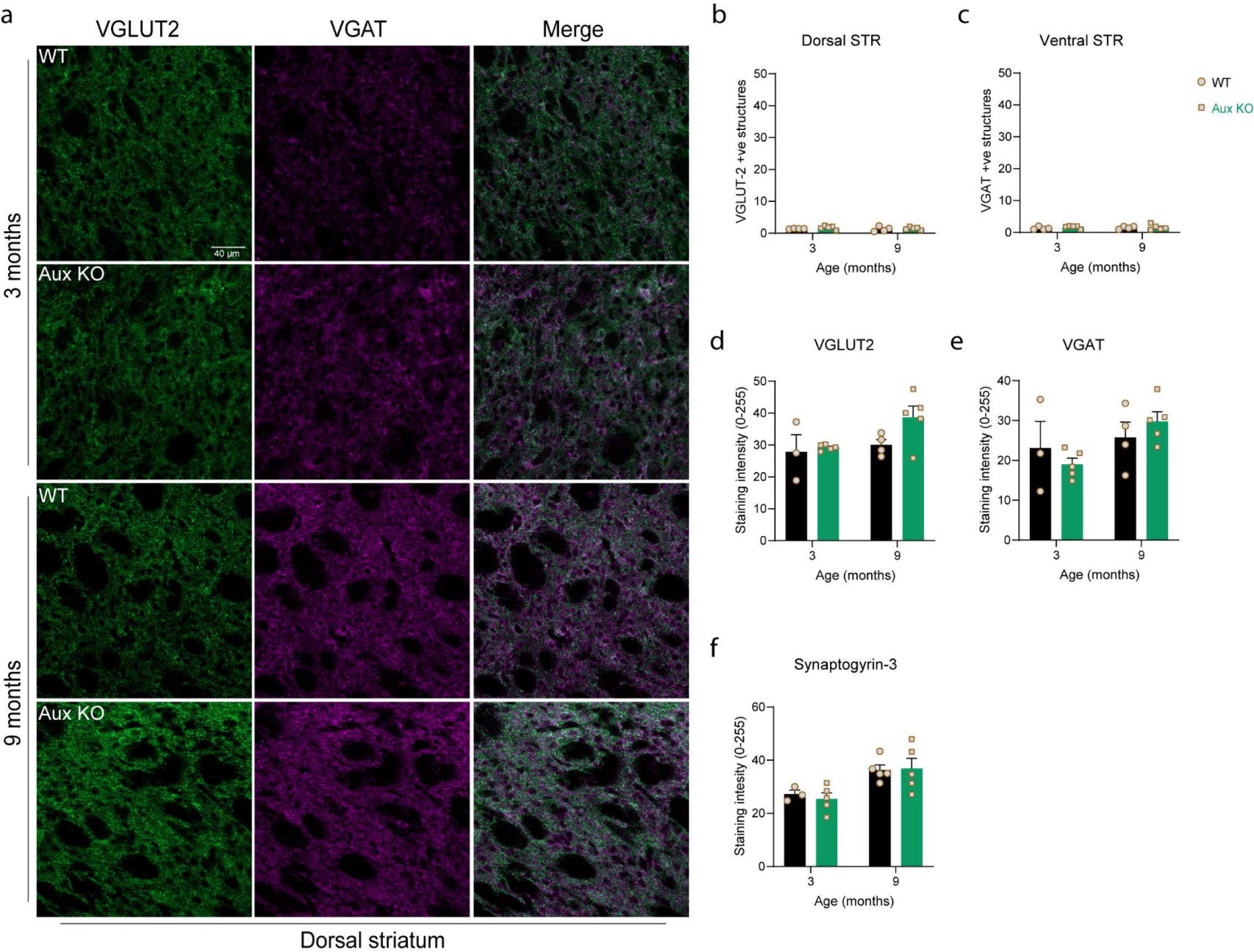
Axonal deformities were not seen in glutamatergic and GABAergic termini. **a.** Representative images showing dorsal striatum immunostained for glutamatergic marker VGLUT2 and GABAergic marker VGAT, at 3 and 9 months of age, in WT and Aux KO mice. Scale bar: 40 μm. **b.** Quantitation for VGLUT2+ve large structures/whirls in dorsal striatum of WT and Aux KO mice. We did not observe any differences between the two genotypes. **c.** Quantitation for VGAT+ve large structures in dorsal striatum of Aux KO mice which revealed no alterations. **d.** Expression of VGLUT2 in the dorsal striatum of WT and Aux KO mice at 3 and 9 months. **e.** Expression of VGAT in the dorsal striatum of WT and Aux KO mice at 3 and 9 months, which did not alter in Aux KOs. **f.** Synaptogyrin-3 expression in the dorsal striatum of WT and Aux KO mice at 3 and 9 months, which was unaltered in Aux KOs (See Figure. 4 for representative images). Statistics: Student’s t-test with Welch’s correction.

**Supplementary Figure. 8:**
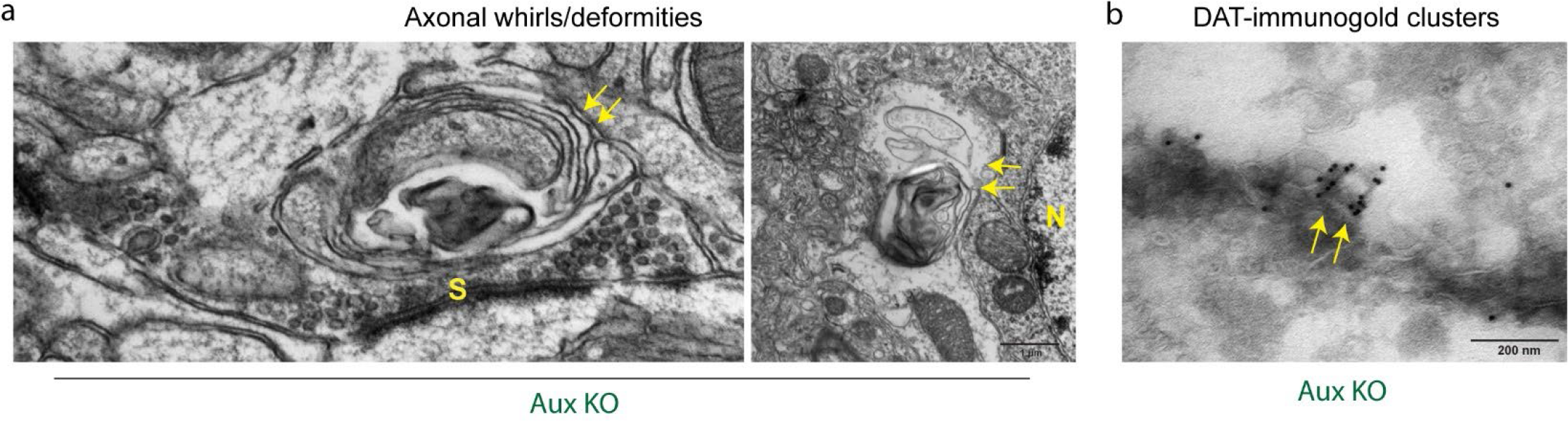
Auxilin KO mice show large axonal whirls/deformities in the dorsal striatum. **a.** Ultrastructure of axonal whirls/deformities in the dorsal striatum of Aux KO mice (arrows). These structures were present both close to synaptic termini (S) and soma (as identified by nucleus, N). Scale bar: 1 μm. **b.** DAT-immunogold clusters in the dorsal striatum of Aux KO mice (arrows). Scale bar: 200 nm.

**Supplementary Figure. 9:**
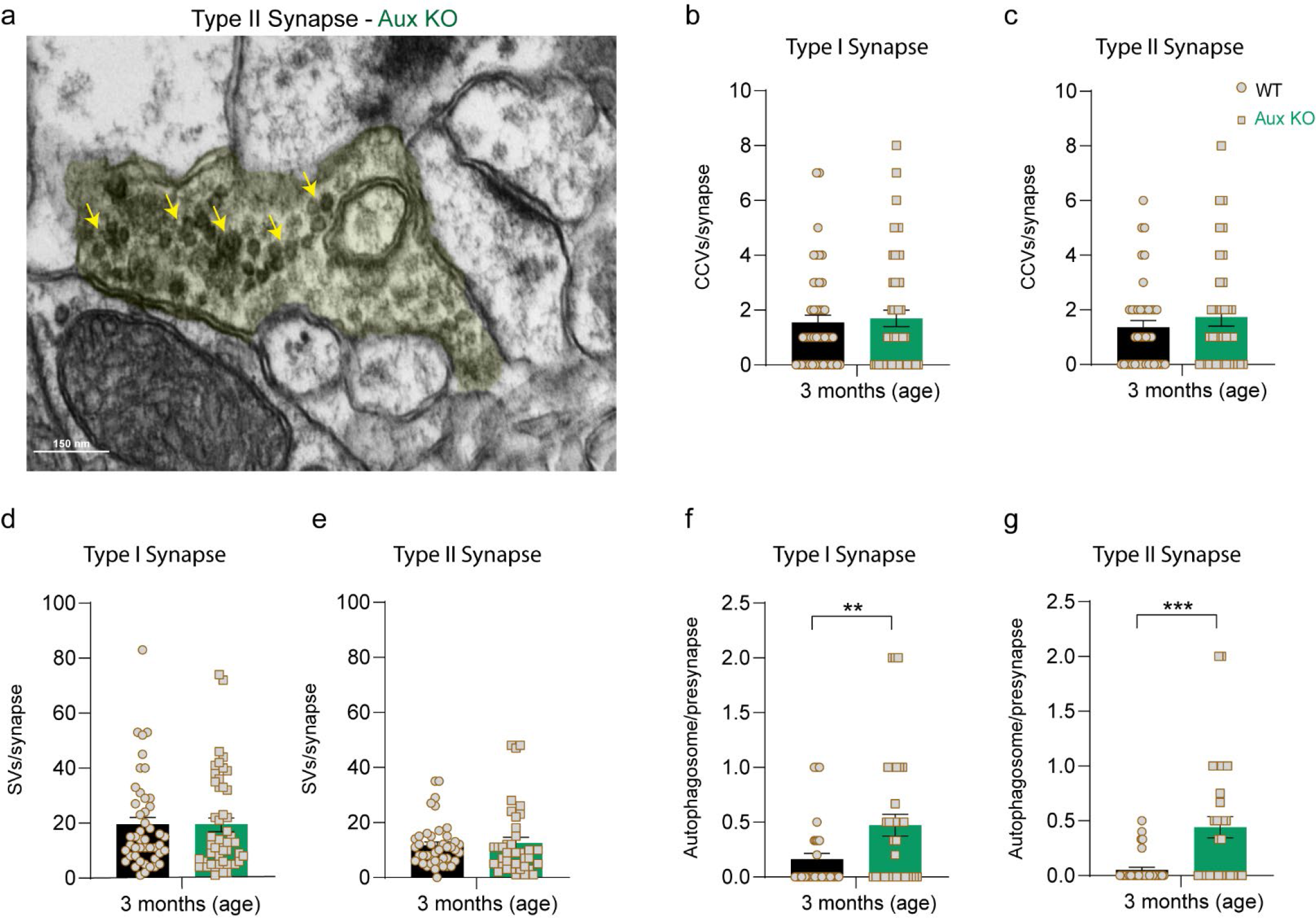
Synaptic autophagy clears accumulated CCVs. **a.** Representative EM image of Aux KO Type I synapse (shaded in yellow) showing CCVs or clathrin cage accumulation, as well as SV clusters (arrows). Scale bar: 150 μm. **b.** CCVs number in Type I synapses, which showed a minor increase in Aux KOs. **c.** CCVs number in Type II synapses, which showed a minor increase in Aux KO mice. **d.** SVs number in Type I synapses, which was not altered. **e.** SVs number in Type II synapses, which also did not change significantly in Aux KOs. **d.** Autophagosomes per presynaptic terminal in Type I synapses of dorsal striatum in WT and Aux KOs, which were significantly higher in Aux KOs. **e.** Autophagosomes per presynaptic terminal in Type II synapses, which were also increased significantly in Aux KO mice. Student’s t-test with Welch’s correction. *\*\*p<0.01, *** p<0.001*

**Supplementary Table 1:**
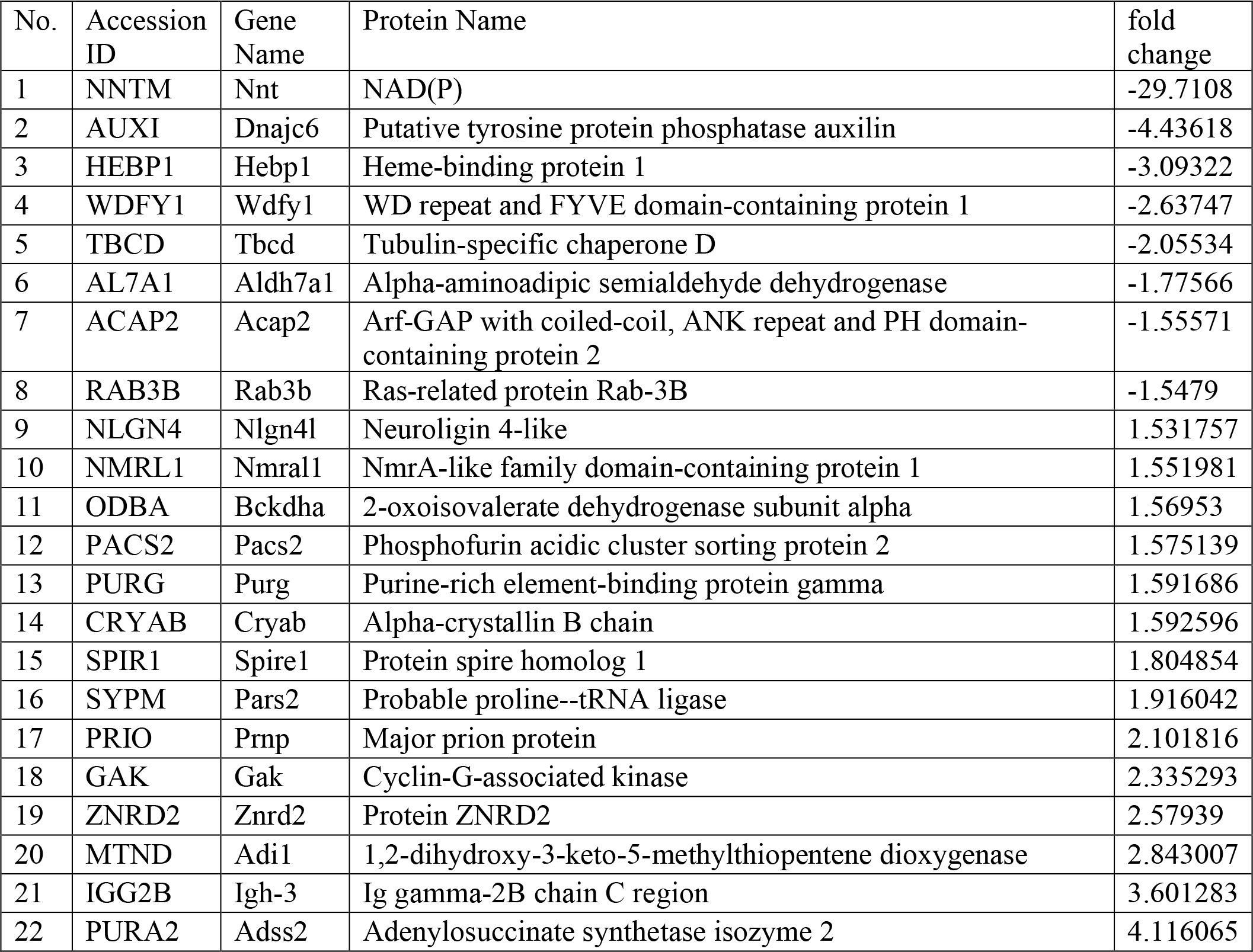
Proteins that are significantly changed in the proteomic analysis of brains of auxilin KO mice, in comparison to WT (age: 3 months).

**Supplementary Table 2:**
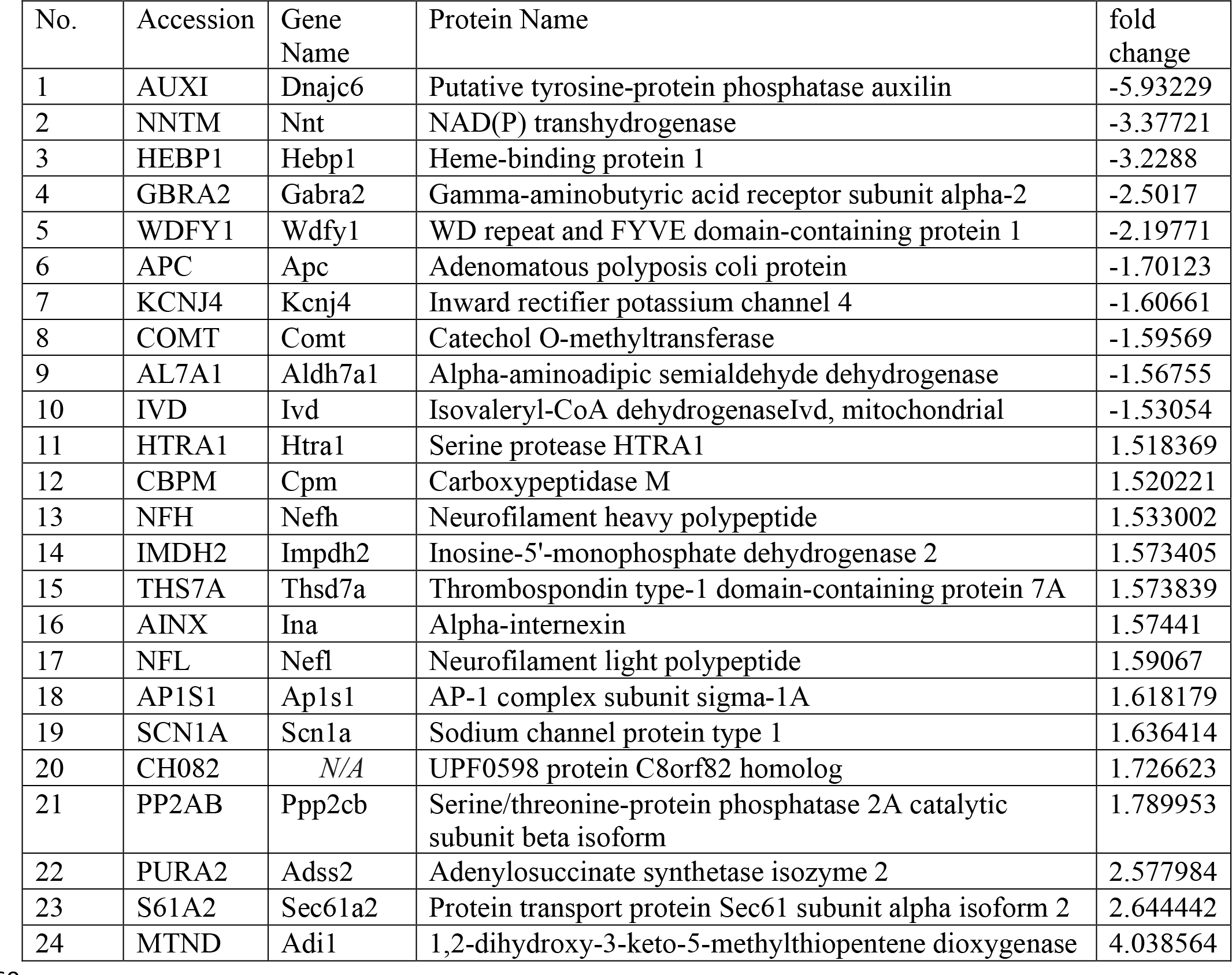
Proteins that are significantly changed in the proteomic analysis of synaptosomes prepared from the brains of auxilin KO mice, in comparison to WT (age: 3 months).

**Supplementary Table 3:**
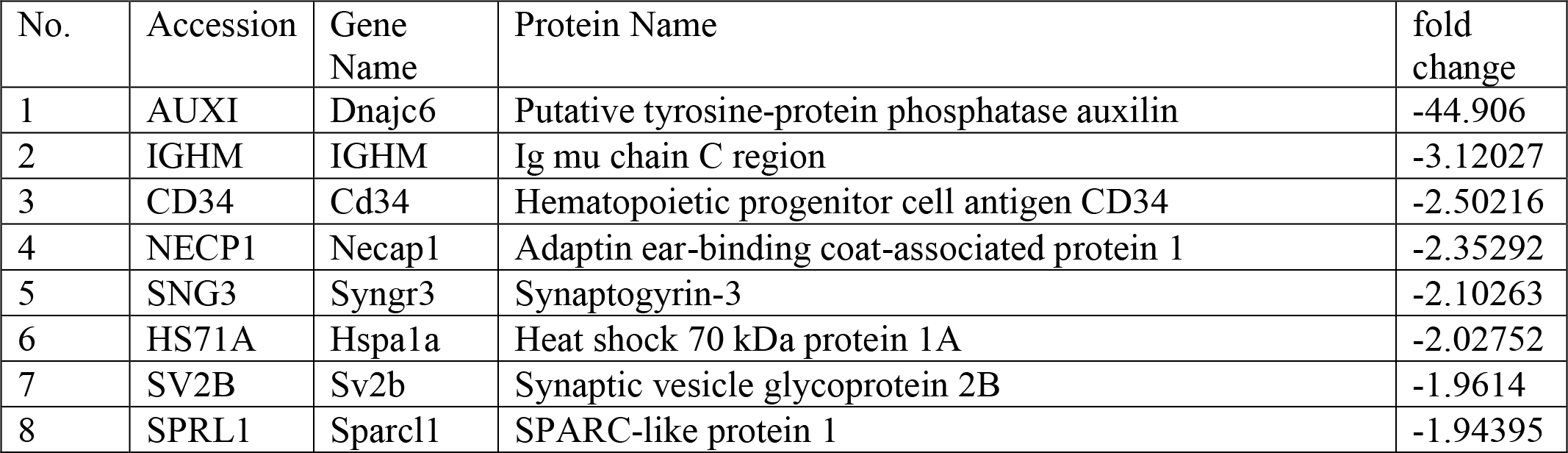
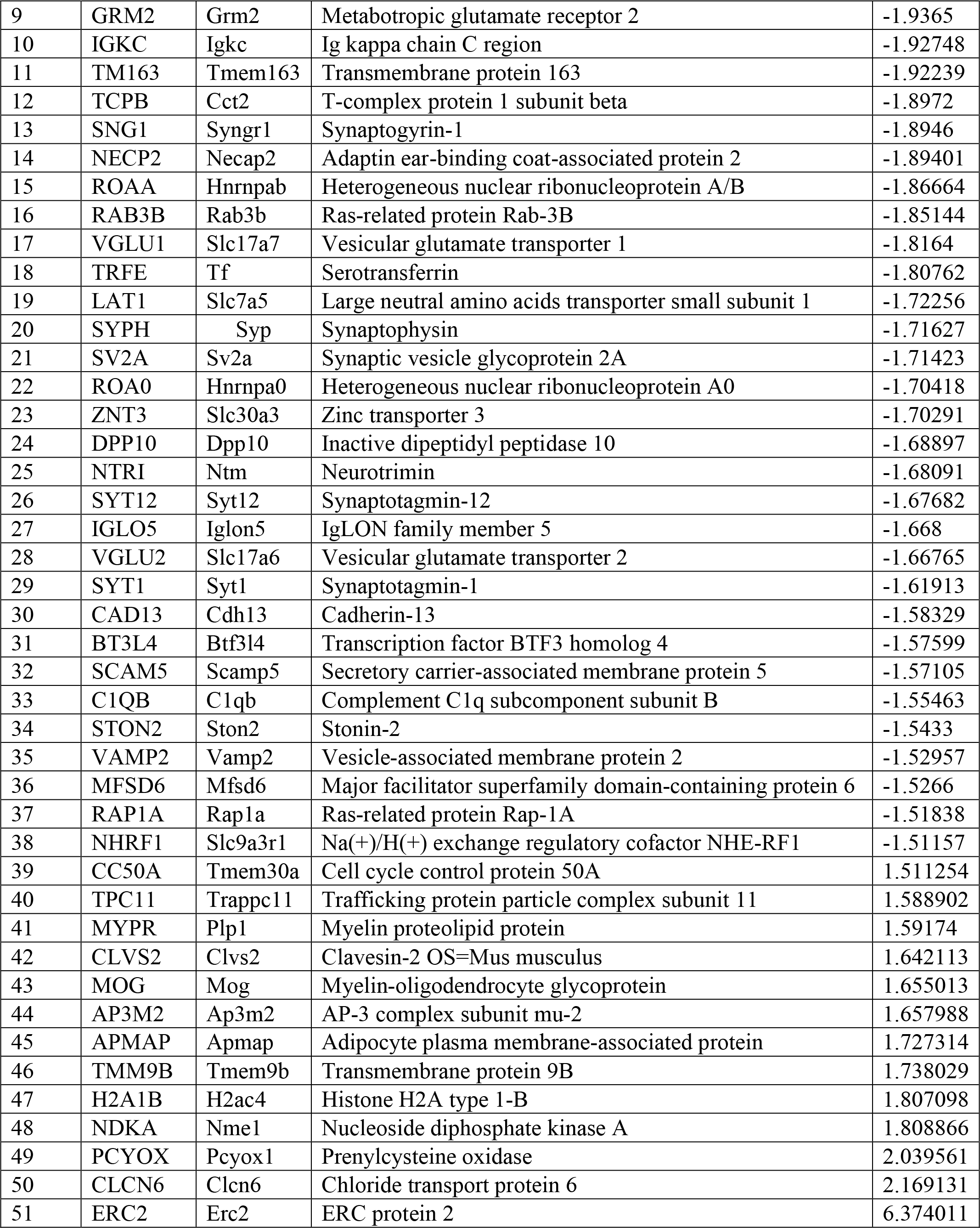
Proteins that are significantly changed in the proteomic analysis of CCVs prepared from the brains of auxilin KO mice, in comparison to WT, at 3 months of age.

**Supplementary Table 4:** Proteins that are significantly changed in the proteomic analysis of synaptosomes prepared from the brains of auxilin KO mice, in comparison to WT, at symptomatic age of 9 months.

| No. | Accession | Gene Name | Protein Name | fold change |
| --- | --- | --- | --- | --- |
| 1 | AUXI | Dnajc6 | Putative tyrosine-protein phosphatase auxilin | -3.82296 |
| 2 | RGS6 | Rgs6 | Regulator of G-protein signaling 6 | -1.66333 |
| 3 | ADA22 | Adam22 | Disintegrin and metalloproteinase domain-containing protein 22 | -1.61876 |
| 4 | ADT1 | Slc25a4 | ADP/ATP translocase 1 | -1.59798 |
| 5 | 2AAB | Ppp2r1b | Serine/threonine-protein phosphatase 2A | -1.57516 |
| 6 | 2AAA | Ppp2r1a | Serine/threonine-protein phosphatase 2A | -1.54334 |
| 7 | PADI2 | Padi2 | Protein-arginine deiminase type-2 | -1.53356 |
| 8 | PRAF3 | Arl6ip5 | PRA1 family protein 3 | -1.52679 |
| 9 | MGLL | Mgll | Monoglyceride lipase | -1.51729 |
| 10 | MTOR | Mtor | Serine/threonine-protein kinase mTOR | -1.50276 |
| 11 | CRIP2 | Crip2 | Cysteine-rich protein 2 | 1.543321 |
| 12 | NFL | Nefl | Neurofilament light polypeptide | 1.545559 |
| 13 | AINX | Ina | Alpha-internexin | 1.557636 |
| 14 | CADH2 | Cdh2 | Cadherin-2 | 2.225312 |
| 15 | ABLM2 | Ablim2 | Actin-binding LIM protein 2 | 1.631285 |
| 16 | ACTN1 | Actn1 | Alpha-actinin-1 | 1.5648206 |
| 17 | GAK | GAK | Cyclin-G-associated kinase | 1.4910263 |

